# Structures of human Malic Enzyme 3

**DOI:** 10.1101/2022.08.25.505315

**Authors:** Tsehai A.J. Grell, Mark Mason, Aaron A. Thompson, Jose Carlos Gómez-Tamayo, Daniel Riley, Michelle V. Wagner, Ruth Steele, Rodrigo F. Ortiz-Meoz, Jay Wadia, Paul L. Shaffer, Gary Tresadern, Sujata Sharma, Xiaodi Yu

## Abstract

Malic enzymes (ME1, ME2, and ME3) are involved in cellular energy regulation, redox homeostasis, and biosynthetic processes, through the production of pyruvate and reducing agent NAD(P)H. Recent studies have implicated the third and least well-characterized isoform, mitochondrial NADP^+^-dependent malic enzyme 3 (ME3), as a therapeutic target for pancreatic cancers. Here, we utilized an integrated structure approach to capture the structures of ME3 in various ligand binding states. ME3 exists as a stable tetramer and its dynamic Domain C is critical for activity. Catalytic assay results reveal that ME3 is a non-allosteric enzyme and does not require modulators for activity while structural analysis suggests that the inner stability of ME3 domain A relative to ME2 disables allostery in ME3. With structural information available for all three malic enzymes, the foundation has been laid to understand the structural and biochemical differences of these enzymes and could aid in the development of specific malic enzyme small molecule drugs.

## Introduction

Malic enzymes are oxidoreductases (oxidative decarboxylases) that catalyze the reversible conversion of (S)-malate to CO_2_ and pyruvate, utilizing NAD^+^ or NADP^+^ as a cofactor and a divalent metal ion (Mg^2+^ or Mn^2+^) for catalysis [1]. There are three known human malic enzymes classified by their sub-cellular localization and cofactor-dependence: cytosolic NADP^+^-dependent isoform 1 (c-NADP^+^-ME, ME1), mitochondrial NADP^+^-dependent enzyme isoform 3 (m-NADP^+^-ME, ME3) and mitochondrial NAD(P)^+^-dependent isoform 2 (m-NAD(P)^+^-ME, ME2) that can use either NAD^+^ or NADP^+^ but prefers NAD^+^ under physiological conditions [2–8].

The structure and function of ME1 and ME2 have been studied in a number of organisms including *Columba livia* [7, 9], *Ascaris suum* [10–14] and *Homo sapiens* [15–20]. ME1 and ME3 are non-allosteric enzymes, but ME2 can be allosterically activated by fumarate and inhibited by ATP [21]. Despite the striking different catalytic mechanisms, both ME1 and ME2 adopt the same homo-tetrameric quaternary structure best described as a dimer of dimers with independently functioning protomers [22]. Each protomer harbors four domains (A, B, C, and D) with the catalytic site located at the cleft between Domains B and C. Open and closed forms involved in cofactor, substrate, and metal ion binding were captured from the crystal structures of ME1 and ME2 [9, 11, 13, 15, 16, 18, 20]. In the open form, the active site is more solvent exposed, allowing the binding of cofactor, substrate, and metal ion. These binding events bring Domain C into close proximity to Domain B, synergistically closing the active site, and inducing the closed form of the enzyme [1]. In addition to the active site, an allosteric site was observed at the dimer interface, formed by residues from Domain A. Fumarate binding at this site results in allosteric activation of ME2 [14, 17, 23]. Although the residues that form the allosteric site are conserved in ME1, allosteric activation has not been observed for this enzyme [20].

Recently, the ME3 isoform has garnered interest due to its link with pancreatic cancer pathology; overexpression of ME3 promotes pancreatic tumor proliferation, invasiveness and metastasis [24]. Targeting ME3 provides a viable strategy to treat a substantial subset of pancreatic ductal adenocarcinoma (PDAC) where the *ME2* gene is deleted [25]. However, there is limited functional and biochemical characterization of ME3, and no structural information is available. Therefore, we sought to obtain essential structural information of ME3 and dissect its reaction mechanism.

## Results

### Crystal structures of human ME3

Purified human ME3 forms a tetramer in solution with a molecular weight of approximately 248 kDa, as indicated by analytical SEC and shows activity with NADP^+^ but not NAD^+^ (Figure S1). The crystal structures of human ME3 were solved here with and without NADP^+^ to 2.49 Å and 1.94 Å resolution, respectively (Figure 1a, Table 1). Both structures revealed two monomers in the asymmetric unit, and a tetrameric unit can be observed via the crystallographic 2-fold symmetry operation (Figure 1a). Both structures revealed high structural similarity with the root mean square (r.m.s.) deviations of 0.25, 0.32, and 0.34 Å between the equivalent monomers, dimers, and tetramers of the two complexes, respectively. Based on the literature precedence of using ME2 sequence numbering, each monomeric subunit contains density for amino acid (a.a.) residues 23/24-584 [9, 20] (Figure S2 and method section for more details).

**Table 1.**
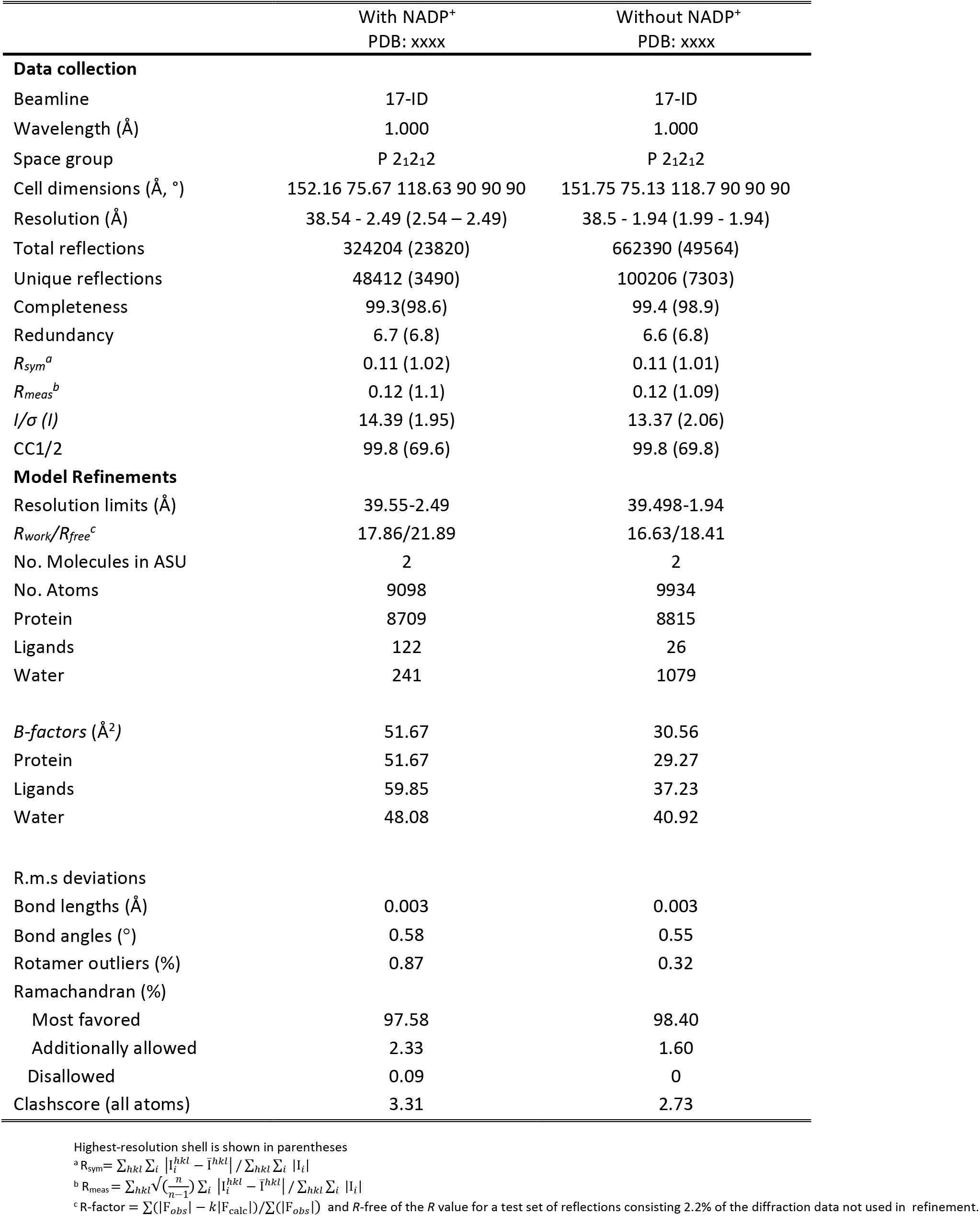
X-ray Data collection, reconstruction, and model refinement statistics.

**Figure 1.**
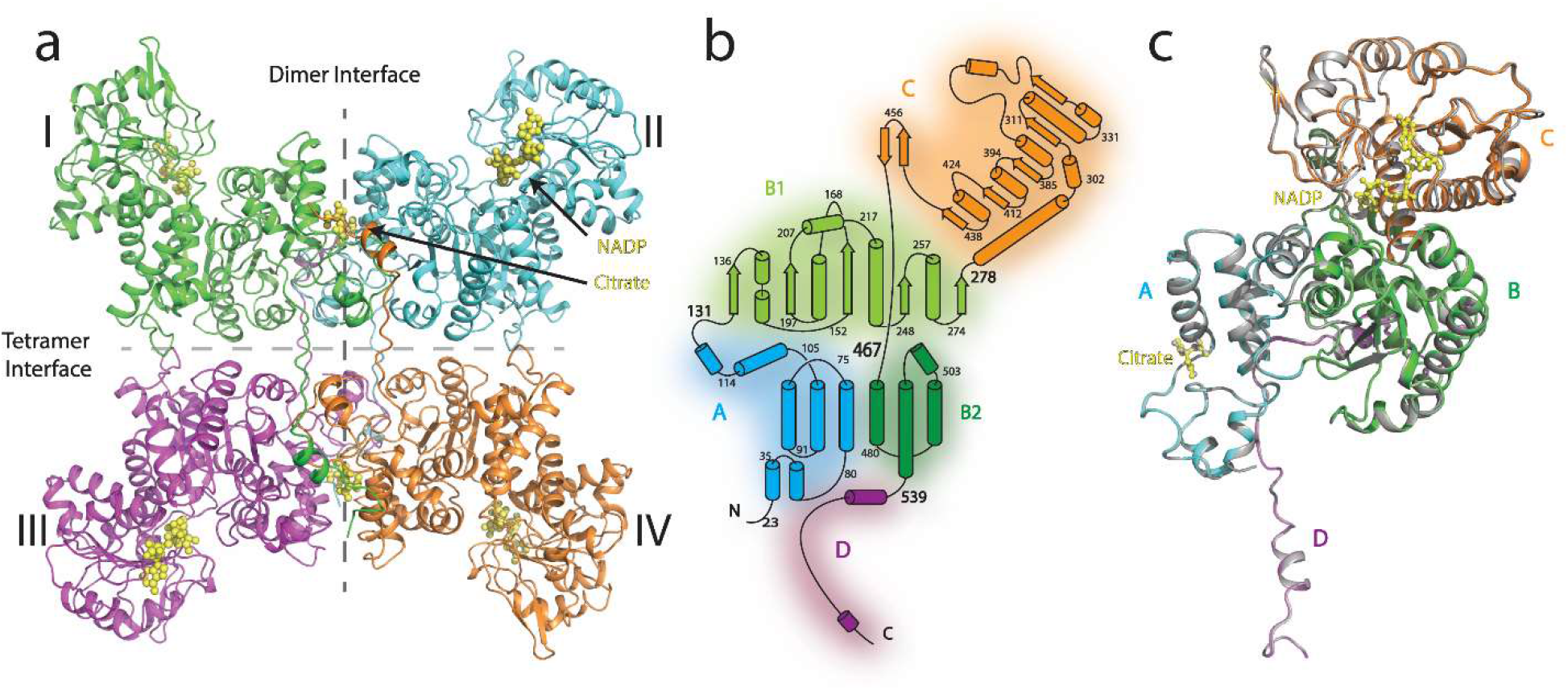
ME3 architecture. (A) The overall architecture of ME3 adopts a dimer of dimer tetrameric assembly with each monomer (protomer) colored in green, blue, pink and orange respectively. The dimeric and tetrameric interfaces are indicated by dashed lines. The NADP^+^ cofactor and the citrate molecule are shown as yellow ball and sticks and indicated by black arrows. (B) Topology diagram of the protomeric subunit of the ME3 tetramer highlighted by domain: Domain A in Blue; Domain B, divided into two subunits B1 and B2, in light and dark Green respectively; Domain C in orange; and Domain D in purple α-helices are represented by cylinders and β-strands arrows. (C) Overlay of Protomers of ME3 structures with NADP^+^ (colored by domain as seen in (B)) and without NADP^+^ (in gray)

Similar to ME1 and ME2, the human ME3 monomer can be organized into four domains [9, 15, 18, 20] (Figure 1b). Domain A (a.a. 23-130) is predominantly helical and is connected to Domain B by Loop AB (a.a. 123-130). Domain B can be split into two subdomains (B1: a.a. 131-277, B2: a.a. 467-538), with Domain C (a.a. 278-466) sandwiched between the subdomains. Domain D (a.a. 539-582) participates in both the tetramer and neighboring dimer formation, functioning as an interdomain linker. Superposition of the two ME3 crystal structures using Domains A and B as the reference reveals a slight opening of the Domain C for the structure without NADP^+^ (Figure 1c). A 1.65° rigid body rotation is needed to fully overlay the two structures.

### Structural comparison of human malic enzymes

The overall fold of human ME3 is highly conserved with the global r.m.s. deviations of 4.02 Å and 6.72 Å when compared to human ME1 and ME2, respectively. Superpositions of pairs of equivalent domains further improves the structural similarities and highlights the presence of subtle domain rearrangements within malic enzyme protomers (Table S1). Noticeable differences are observed between Domains B and C of the protomers, corresponding to the open and closed forms. Superposition, using Domains A and B as the reference, reveals small variations of the ME3 structures compared with the open form structures, whilst a more substantial 8° rigid body rotation of Domain C is needed to fully overlay the ME3 structures and the closed form structure (Figure 2a and b, Table S2). Taken together, both the ME3 crystal structures, with and without NADP^+^, are in the open form.

**Figure 2.**
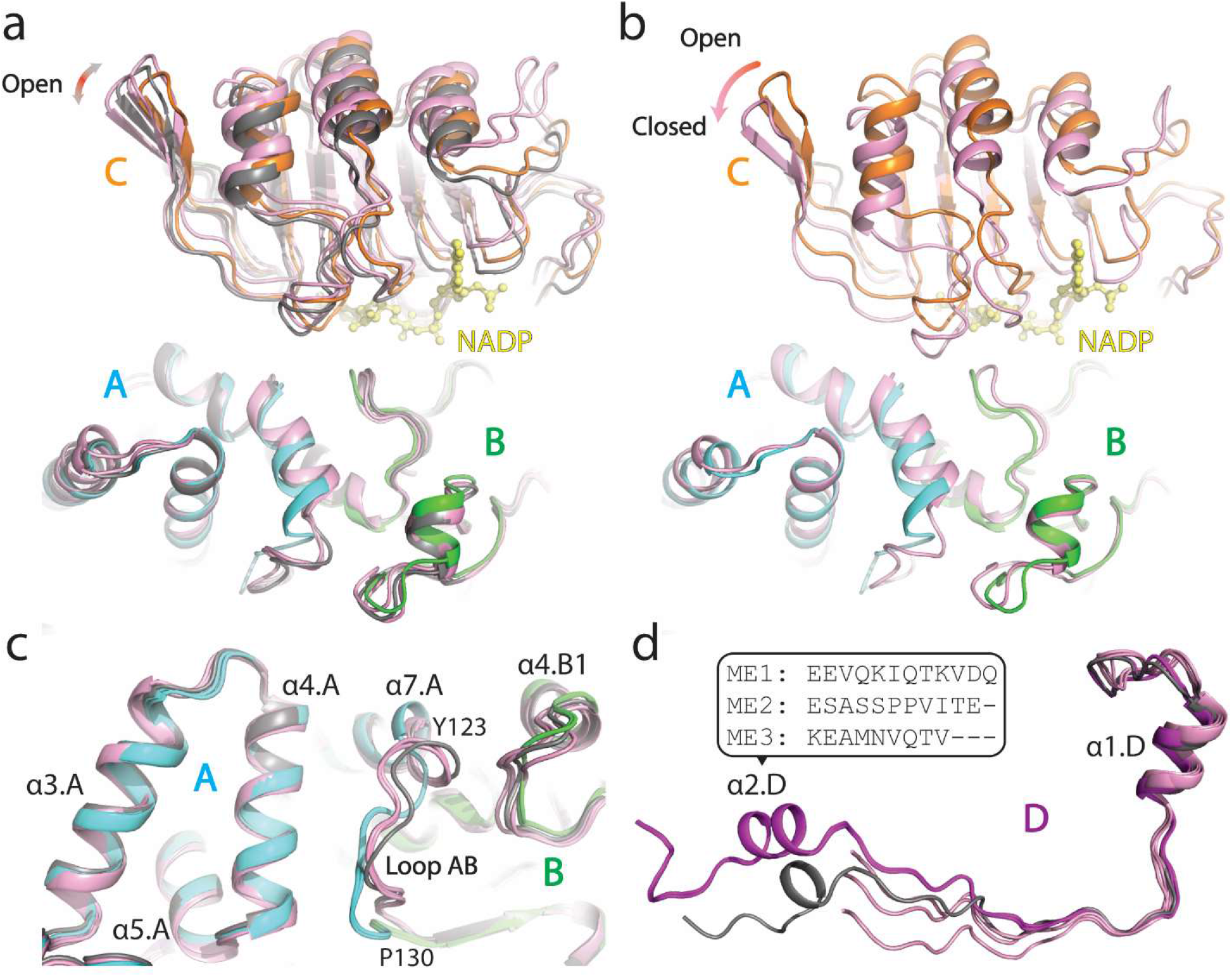
Domain comparison of ME3 structures to open and closed structures of ME2 and ME1. ME3 NADP^+^-bound structure overlaid with Domains A-C of the (A) open forms of ME1 (PDB ID: 3WJA) colored in gray and ME2 (PDB IDs: 1QR6, 1PJL) colored in pink and (B) closed form of ME2 (PDB ID: 1PJ3) colored in light pink. (C) Different orientation of the overlay of Domains A and B of ME3 structures in the presence and absence of NADP^+^ with that of ME1 (PDB ID: 3WJA, gray) and ME2 (PDB IDs: 1QR6, 1PJL, pink). (D) Superposition of Domain D of both ME3 structures (purple), ME1 (PDB ID: 3WJA, gray) and ME2 (PDB IDs: 1QR6, 1PJL, pink). In all panels, ME3 is colored by domain: Domain A in Blue; Domain B in Green; Domain C in orange; and Domain D in purple.

Large degrees of structural heterogeneities were observed among the three malic enzyme isoforms at the Loop AB (Figure 2c). Another noticeable difference was observed in Domain D (Figure 2d) where ME1 and ME3 structures adopt an additional α-helix that interacts with the neighboring dimer and potentially enhances tetramer formation; whilst this corresponding α-helix was less resolved in the ME2 structures. Noticeable rigid body movements were also observed in the tetramer organization when superposition is performed using one protomer as the reference (Figure S3a). These observations indicate the three malic enzyme isoforms share conserved structural architectures with subtle global and local variations.

### Determinants of cofactor selectivity in ME3

One NADP^+^ cofactor was modelled between Domains C and B of ME3 protomers from crystals soaked in the presence of NADP^+^ (Figure S4a). The cofactor interacts almost exclusively with residues from Domain C, with residues from Domain B helping to position the alpha and beta phosphates and the primary amide of the nicotinamide group (Figure 3a). Like ME1 (PDB ID: 1GQ2), the NADP^+^ phosphate group appended to the ribose 2′OH is accommodated by a conformational shift in the β2-β2′ hairpin proximal to this group when compared to the NAD bound ME2 structure (PDB ID: 1PJ3) (Figures 3a and S5). The 2′OH of NADP^+^ is positioned to interact with Ser346, which is a Lys residue in the case of ME2 and would clash with the phosphate group without an alternate rotamer configuration. In the case of ME3, the Lys register is shifted by one residue thereby positioning Lys347 for an interaction with the phosphate group (Figure 3a). Although this Lys residue is conserved in ME1, it does not interact with NADP^+^ (Figure S5c, S2) and mutation of this residue has less than a twofold effect on *Km* of ME1 for NADP^+^, and no significant effect on the *Kcat* of the enzyme [26]. Further work is needed to ascertain if Lys347 is involved in NADP^+^ selectivity or catalysis of ME3. Another difference between the NAD^+^ and NADP^+^-dependent malic enzymes is at position 362. In ME3 and ME1 this residue is a lysine which further stabilizes the phosphate group and selects for NADP^+^. ME2 has a glutamine residue at this position (Figure 3a, S5), and when mutated to a lysine the cofactor selectivity of ME2 strictly switches to NADP^+^ [27].

**Figure 3.**
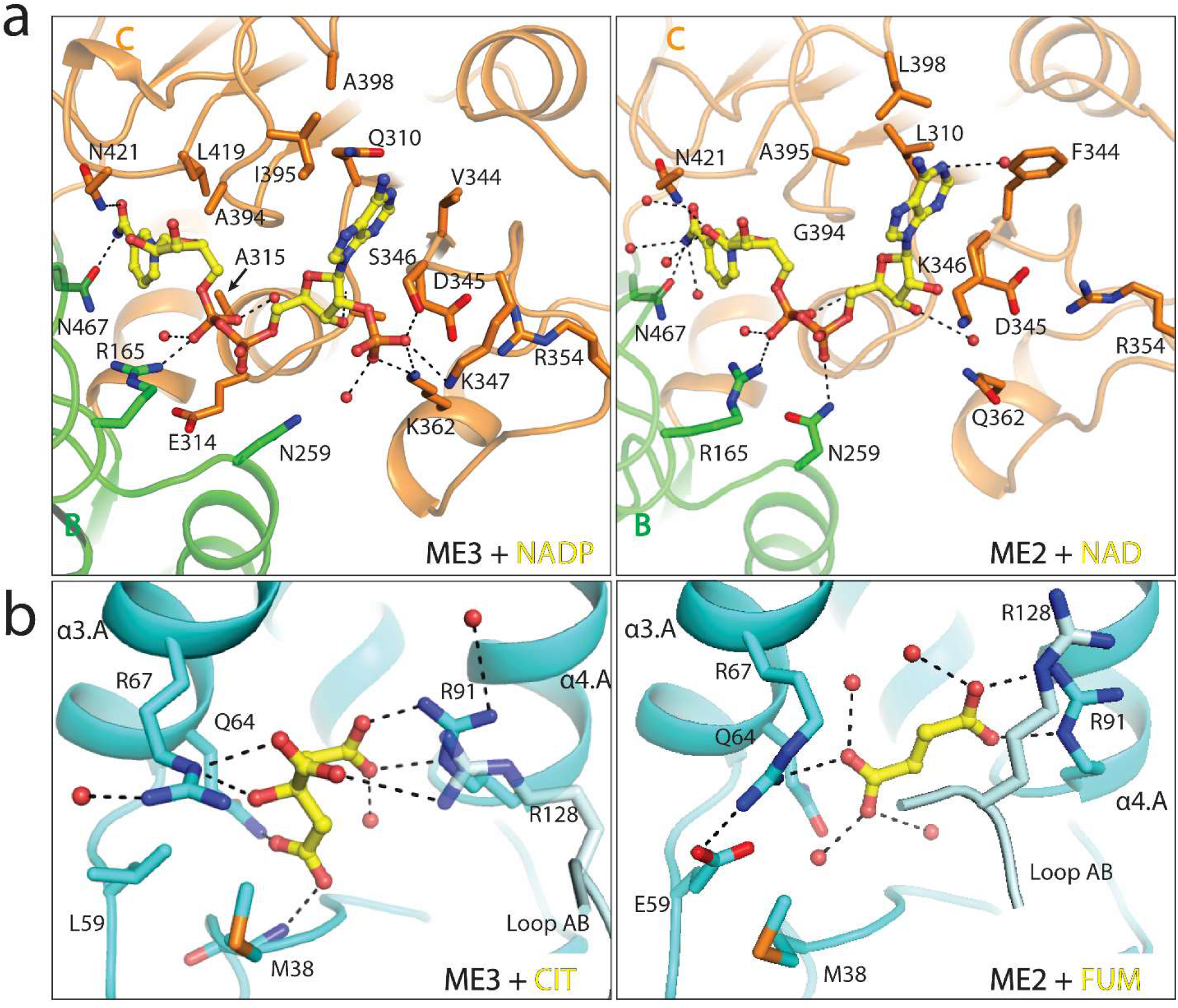
Comparison of cofactor and ligand binding in mitochondrial Malic enzyme. (A) NADP^+^/ NAD cofactors (yellow ball-and-sticks) bind to the in the active sites of ME3 (left) and ME2 (right), respectively. NAD(P) binding is mediated primarily by residues of domain C (orange) and a few interactions with domain B1 (green). (B) Citrate (yellow ball-and-sticks) binds to the putative fumarate binding site in ME3 (left) in a similar manner to that of fumarate (yellow ball-and-sticks) in ME2 (right). This allosteric binding site is located at the malic enzyme dimer interface and is created by residues (sticks) in Domain A (colored in blue) from one protomer and residues from loop AB from the dimeric protomer (colored light cyan). Of note, citrate forms ionic interactions with residues at positions 38, and 128 of the neighboring dimer which are not observed in the fumarate-bound ME2 structures.

### Cryo-EM structures of human ME3

Cryo-EM structures of human ME3 in the apo state as well as with additional NADP^+^ were determined at resolutions comparable to the crystal structures (Figure 4a, S6 and Table 2). The EM structures form a tetrameric complex with D2 symmetry. 3D classifications showed predominantly a tetrameric complex, indicating that ME3 forms a stable tetramer complex in solution. Superposition of the crystal and cryo-EM structures of ME3 revealed a conserved overall topology, though subtle differences were observed (Figure S3b). The apo EM structure exhibits a more open conformation with the active site more exposed to solvent, than the apo crystal structure (Figure 4b). Superposition of NADP^+^-bound EM and crystal structures revealed high similarity with the global r.m.s. deviation around 1.17 Å, indicating cofactor binding rigidifies the pose of ME3 (Figure 4b, S4b). Loop AB has decreased local resolution in all EM structures (Figure 4c). Molecular dynamic (MD) simulations using Rosetta [28] allowed the rebuilding of the loop regions and presumably indicate that the Loop AB can adopt various conformations (Figure S7). In addition, loop 564-567 of Domain D forms additional hydrophobic and main chain hydrogen bond interactions with Domain A from its own protomer in the cryo-EM structure, potentially enhances the tetramer formation (Figure 4d).

**Table 2.**
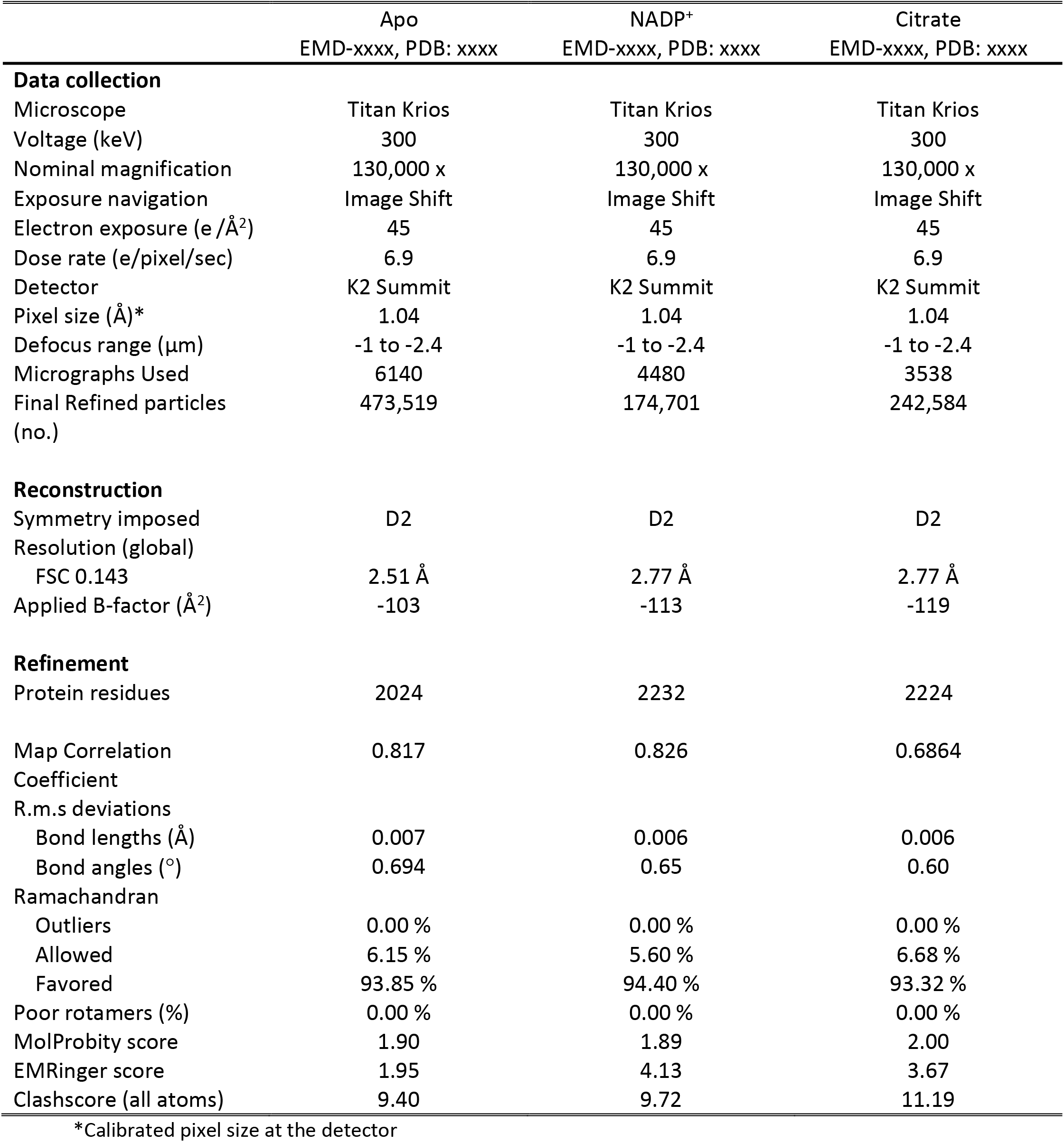
Data collection, reconstruction, and model refinement statistics.

|  | Apo<br>EMD-xxxx, PDB: xxxx | NADP<sup>+</sup><br>EMD-xxxx, PDB: xxxx | Citrate<br>EMD-xxxx, PDB: xxxx |
| --- | --- | --- | --- |
| **Data collection** |  |  |  |
| Microscope | Titan Krios | Titan Krios | Titan Krios |
| Voltage (keV) | 300 | 300 | 300 |
| Nominal magnification | 130,000 x | 130,000 x | 130,000 x |
| Exposure navigation | Image Shift | Image Shift | Image Shift |
| Electron exposure (e /Å<sup>2</sup>) | 45 | 45 | 45 |
| Dose rate (e/pixel/sec) | 6.9 | 6.9 | 6.9 |
| Detector | K2 Summit | K2 Summit | K2 Summit |
| Pixel size (Å)* | 1.04 | 1.04 | 1.04 |
| Defocus range (μm) | -1 to -2.4 | -1 to -2.4 | -1 to -2.4 |
| Micrographs Used | 6140 | 4480 | 3538 |
| Final Refined particles<br>(no.) | 473,519 | 174,701 | 242,584 |
| **Reconstruction** |  |  |  |
| Symmetry imposed | D2 | D2 | D2 |
| Resolution (global) |  |  |  |
| FSC 0.143 | 2.51 Å | 2.77 Å | 2.77 Å |
| Applied B-factor (Å<sup>2</sup>) | -103 | -113 | -119 |
| **Refinement** |  |  |  |
| Protein residues | 2024 | 2232 | 2224 |
| Map Correlation<br>Coefficient | 0.817 | 0.826 | 0.6864 |
| R.m.s deviations |  |  |  |
| Bond lengths (Å) | 0.007 | 0.006 | 0.006 |
| Bond angles (°) | 0.694 | 0.65 | 0.60 |
| Ramachandran |  |  |  |
| Outliers | 0.00 % | 0.00 % | 0.00 % |
| Allowed | 6.15 % | 5.60 % | 6.68 % |
| Favored | 93.85 % | 94.40 % | 93.32 % |
| Poor rotamers (%) | 0.00 % | 0.00 % | 0.00 % |
| MolProbity score | 1.90 | 1.89 | 2.00 |
| EMRinger score | 1.95 | 4.13 | 3.67 |
| Clashscore (all atoms) | 9.40 | 9.72 | 11.19 |

\*Calibrated pixel size at the detector

**Figure 4:**
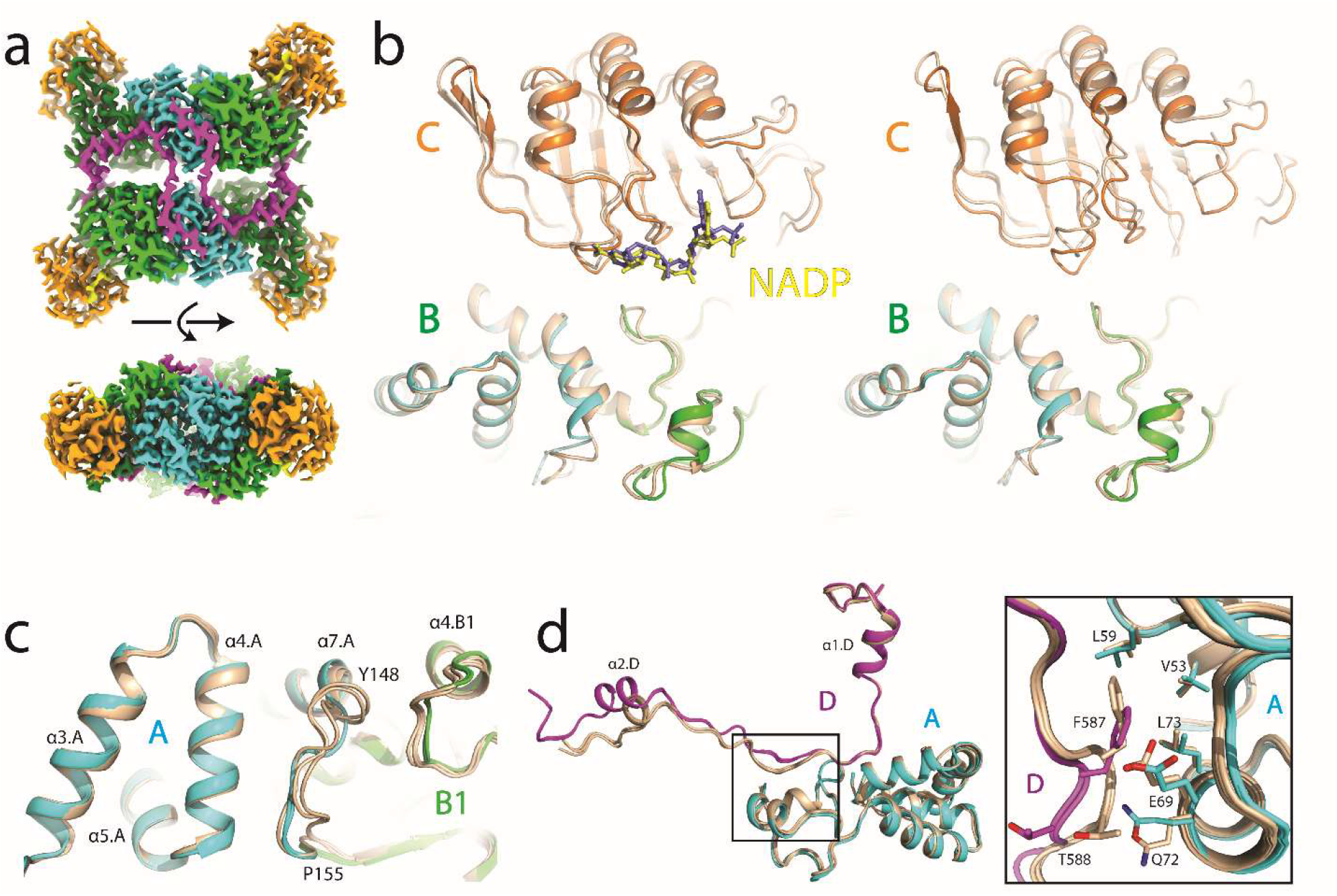
Cryo-EM structures of ME3. (A) Side and top-down views of ME3 cryo-EM density map showing with each protomer colored by domains. (B) Comparisons of the active site (Domains B and C) of ME3 cryo-EM structures and the corresponding X-ray structures. (Top panel) Overlay of NADP^+^-bound cryo-EM and the X-ray structures with the NADP^+^ cofactor represented as sticks and colored yellow and blue, respectively. (Bottom panel) Superposition citrate-bound cryo-EM and X-ray structures of ME3. (C) Overlay of Domain A and D from NADP^+^-and citrate-bound cryo-EM and X-ray structures. Zoomed in view of the interactions between the two domains (A and D) are shown in the inset. (D) Overlay of Domain A and B from a protomer of NADP^+^-and citrate-bound cryo-EM and X-ray structures highlighting the dynamic features of loop AB (Y123-P130). For panels B-D, the cryo-EM structures are colored in wheat and the X-ray structures are colored by domain: Domain A in Blue; Domain B1 and B2, in light and dark green respectively; Domain C in orange; and Domain D in purple.

### EM analysis reveals dynamics in malic enzyme holoenzyme formation

To investigate the dynamics of individual protomers in various ligand bound states, the final reconstructed EM maps were used to perform symmetry expansion followed by signal subtraction on the individual protomers (Figure S8a). 3D classifications without alignment or symmetry operations showed the NADP^+^-bound state is more conformationally homogeneous, whereas the apo state is conformationally heterogeneous with the most noticeable flexibilities at domain C (Figure S8b-e). Multi-body refinement with two masks, corresponding to Domains A+B and C, further revealed that Domain C of the apo state is dynamic, spanning widely open and closed conformations. NADP^+^ binding to domain C limits its movements. Interestingly, a short opening movement was also observed in the NADP^+^-bound state which may be due to the lack of full occupancy of the NADP^+^ site, representing the cofactor prior capturing state or post releasing state (video 1).

### ME3 is a non-allosteric enzyme

Non-protein density was identified at the putative allosteric site of both ME3 crystal structures and was modelled as a citrate molecule based on the shape of the electron density and components of the crystallization conditions (Figure S4c and d). From the ME3 crystal structures, the citrate forms interactions with the conserved residues Arg67, Arg91 and Gln64, which are involved in fumarate binding in ME2 structures (Figure 3b, S2). One notable difference is observed at position 59. In ME2, Glu59 forms a salt bridge with Arg67 and is critical for the allosteric activation of ME2 [31]. In ME3 this corresponding residue is changed to a leucine and Arg67 adopts an alternate rotamer pointing towards the dimeric protomer. Since human ME3 crystals formed only in the presence of citrate, single-particle cryo-EM was used as a complimentary technique to investigate ME3 structural plasticity and the possible structural effect of citrate binding. Superposition of apo and citric bound EM structures revealed high similarity with the global r.m.s. deviation around 0.48 Å, indicating citrate has no apparent structural impact on ME3. Interestingly, the citrate signal was not identified at the putative allosteric site in the citrate-bound EM map. One explanation is citrate is a highly negatively charged molecule that exhibits more pronounced effects of radiation damage in structures solved by cryo-EM. Another possibility is an increased dynamic nature of Loop AB at the allosteric site in the EM experiment. The structural plasticity and diversity of the allosteric site indicates specific functional roles for ME2 and ME3, respectively.

To determine whether the presence of citrate or fumarate perturbs the catalytic rates of ME3 and ME2, the formation of NAD(P)H was followed by measuring the increase in absorbance at 340nm at saturating substrate conditions. Consistent with previous studies [29], we observed an increase in ME2 catalytic activity in the presence of 5mM fumarate (but not 5mM citrate) (Figure 5A Left). While a thermal shift assay indicated that citrate, but not fumarate, increased the stability of ME3 complex (Figure S4e), we detect no significant change in the catalytic activity of ME3 in the presence of either 5mM citrate or fumarate (Figure 5A Right). Taken together, these data indicate that ME3 permits allosteric binders, but is not allosterically modulated by citrate or fumarate.

**Figure 5:**
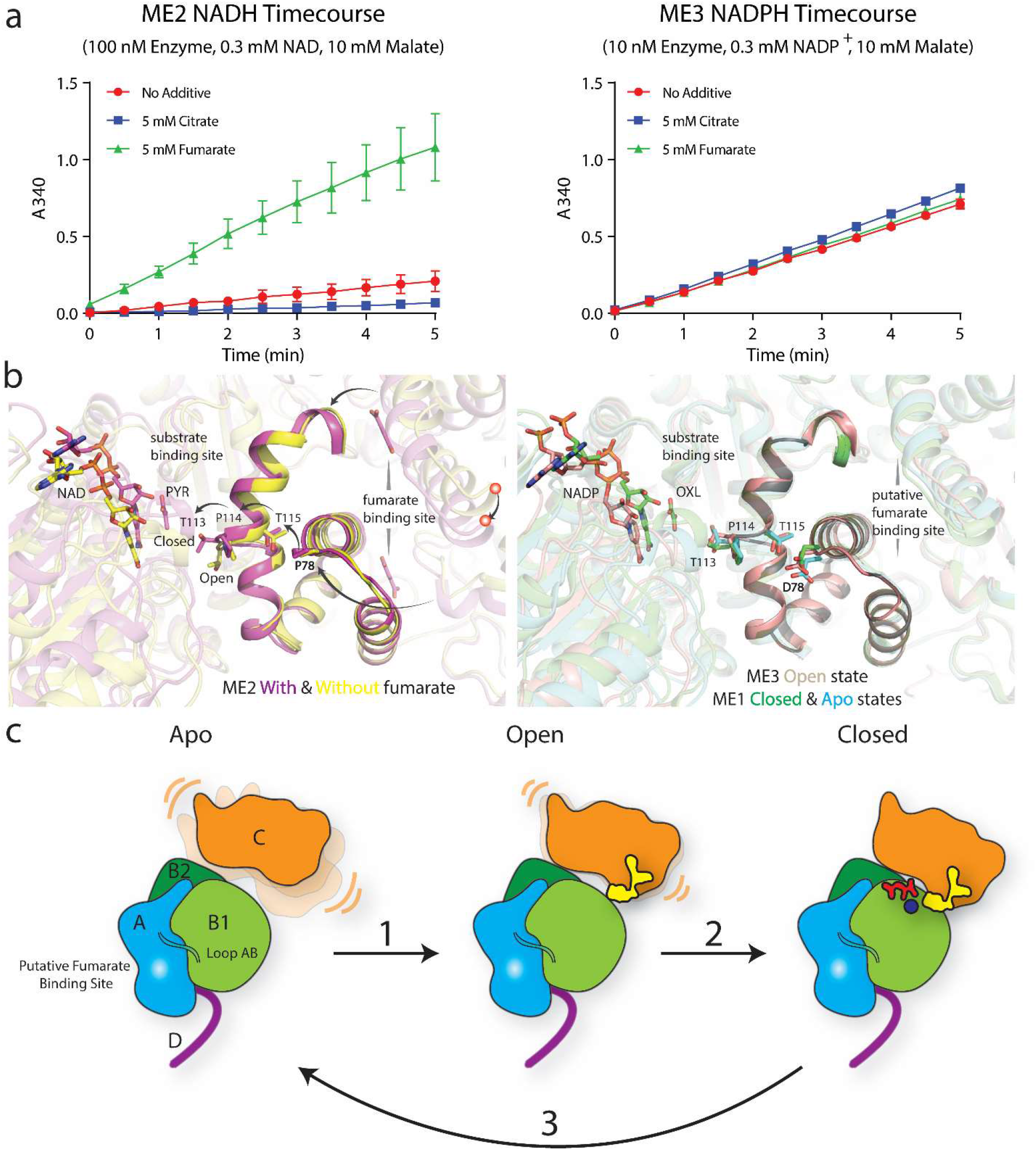
ME3 is a non-allosteric enzyme. **(A)** Malic enzyme biochemistry. Activity assays for ME2 (**Left**) and ME3 (**Right**) performed in the presences of either 5 mM Citrate or Fumarate at saturating conditions. **(B)** Force transition from the fumarate allosteric binding site to the substrate binding site. **Left**, structural overlay of ME2 closed and open states. Red balls highlight the shift of the dimeric protomer upon fumarate binding. Dark arrows represent the path of force transition. **Right**, structural overlay of ME3 open with ME1 closed and apo states. **(C)** Schematic representation of the dynamic state of ligand bound states of ME3. In the apo state, domain C of ME3 is highly flexible and occupies different orientations. Binding of the cofactor NADP^+^ to domain C and a lesser extent to domain B1 stabilizes domain C in an open form. Finally binding of the substrate malate and the catalytic metal is expected to restrict movement of domain C locking the protein into a closed form ready for catalysis.

### Structural analysis reveals, unlike ME2, ME3 is a non-allosteric enzyme

To provide a structural framework for understanding allosteric modulation mechanism for ME2 and ME3, we first focus on the allosteric binding site. The site is composed of Domain A and Loop AB from the dimeric protomer that shield it from the solvent. The fumarate binding to the ME2 allosteric binding site shifts the oligomerization state to tetramer and cooperatively enhances its enzyme activity [22, 29]. Mutations introduced at the ME2 allosteric binding site not only disrupt the oligomerization state, but also silence the modulatory activity of fumarate [21]. In contrast, the same allosteric regulation was not observed for ME1 or ME3 [22].

ME2 structures with and without fumarate have been determined previously (Table S2). Upon fumarate binding, Domain A undergoes a conformational change leading to a closer arrangement of the dimeric interface. The distances measured between the two Thr76 residues at the interface of the two monomers are 11.0 and 14.2 Å for ME2 with and without fumarate respectively. Furthermore, the angle between the helices at the interface is 48.0 and 55.3 degrees for ME2 with and without fumarate structures, respectively (Figure S9). In addition, Phe68 adopts an alternative rotamer that flips inward to fill the empty volume and opens the allosteric pocket that permits fumarate binding (Figure S10) [15, 29]. The binding energy between the two monomers was assessed (prodigy web server [30, 31]) and calculated as −13.8 and −13.3 kcal/mol for ME2 with and without fumarate structures, respectively. The fumarate engagement further stabilizes the ME2 dimer by ^~^6 kcal/mol. Hence, fumarate may regulate the activity of ME2 by stabilizing the closed dimeric interface.

Instead, ME3 intrinsically has increased stability. The calculated binding energy between the ME3 two monomers was stronger, −15.3 kcal/mol. Also, the equivalent distance and angle between monomers are similar to ME2 bound to fumarate (11.2 Å and 49.9 degrees respectively), suggesting ME3 possesses a constitutively more stable and packed dimeric interface. Larger hydrophobic residues (Tyr72 and Phe107 in ME3 instead of Leu72 and Leu107 in ME2, respectively) at the Domain A form a tighter hydrophobic packing which shows reduced flexibility compared to ME2 as observed in the structures described in this work (Figure S13). Additionally, the non-interacting pair Lys73-Ser106 in ME2, is replaced by a strong ionic interaction between Glu73 and Lys106 (Figure S11).

We further investigated how the regulation is transmitted from the allosteric site to the substrate binding site by comparing the difference between ME3 and ME2 structures. Upon fumarate binding in ME2, the Domain A helix II (a.a. 78-91) moves and performs a mechanical push via Pro78 towards Thr115 in the transversal helix (a.a. 103-127) (Figure 5b Left, S12, S13). This push increases the kink formed by Thr113 and Pro114 causing Thr113 to move and face the substrate in the binding site. In contrast, ME3 (also ME1) possesses a similar kink and Thr115 poses as observed in ME2-closed orientation, regardless of the ligand binding (Figure 5b Right, S12, S13) state. Moreover, ME2-Pro78 is replaced with Asp78 in ME3 (also in ME1), that could remove the ability to perform the initial step of the mechanism connecting the allosteric and substrate sites in ME3. Taken together, the analysis suggests that unlike ME2, ME3 is a non-allosteric enzyme.

## Discussion

Here we provide the first atomic structures of human ME3, completing the initial structural characterization for this family of enzymes. Although the overall architecture is conserved for these enzymes, differences were noted in tetramer compactness and domain orientations of the malic enzymes [1]. The ME3 structures in this study adopt an open form, with the apo and citrate-bound EM structures exhibiting an extended open form. The cryo-EM analysis reveals the dynamic nature of malic enzymes: Domain C undergoes large conformational changes upon cofactor binding (Figure 5c and video1). In the apo form, Domain C is highly flexible and the binding of the cofactor NADP+ reduces the conformational flexibility, preforming the active site for metal and substrate binding (Figure 5c). This conformational flexibility was not observed in the crystal structures and highlights the strengths of an integrated structural biology approach. Loop AB was observed to be highly flexible, adopting variable conformations in malic enzyme isoforms. Loop AB sits at the dimer interface and may play a role in allosteric activation of ME2 and dimer compactness in the malic enzymes.

All ME isoforms have a similar quaternary structure with four functional independent protomers ([1] and this study). Both ME1 and ME3 can form a stable tetrameric complex and perturbing the oligomerization state of ME1 does not affect its kinetic properties [22]. While both the oligomerization state and activity of ME2 are closely regulated by the binding of an allosteric modulator, fumarate, at the dimer interface. The fumarate binding may help to mature domain A via tightening the dimeric interface and ultimately trigger the flip of ME2-Thr113 side chain that favors the interaction with the substrate. In contrast, the Thr113 from ME1 or ME3 naturally adopts the favourable pose for substrate binding, hence the cooperation of substrate binding, the putative allosteric binding or oligomerization state was not pronounced. Interestingly, ME1 and ME3 are non-allosteric regulated enzymes but still exist as a stable tetramer and preserve a putative allosteric binding site that allowing allosteric binders, such as citrate (this study). In addition, both ME2 and ME3 play redundant roles in the mitochondrion with striking different catalytic profiles. Further studies are needed to uncover the biological signatures of ME isoforms.

The most recent breakthrough in the structure prediction via computational artificial intelligence approaches allows us to compare the determined structures with the virtual structure at first hand. The AlphaFold2 predicted ME3 model closely resembles the determined structures with notable differences observed in Domains C, D and Loop AB (residues around 125) (Figure S14). The differences may be induced by the ligand binding or the tetrameric complex assembly. We believe AlphaFold2 can generate valuable models to validate experimental results, meanwhile providing structural insights for targets that are non-trivial to be obtained by traditional methods.

The present work highlights an integrative approach utilizing complementary techniques to provide comprehensive structural understanding. The structural information provided in this study could not be obtained by a standalone structural technique. As high resolution cryo-EM technology evolves we expect to tackle broader target classes and with less limitation of size. Also, the combined approaches such as the one highlighted in this study will be leveraged more routinely to gain better functional understanding of a given target and potentially aid rational drug design.

## Acknowledgement

This research used resources of the Advanced Photon Source, a U.S. Department of Energy (DOE) Office of Science User Facility operated for the DOE Office of Science by Argonne National Laboratory under Contract No. DE-AC02-06CH11357. Use of the IMCA-CAT beamline 17-ID at the Advanced Photon Source was supported by the companies of the Industrial Macromolecular Crystallography Association through a contract with Hauptman-Woodward Medical Research Institute. The cryo-EM data were collected at NanoImaging Services (San Diego, CA).

## Author contributions

Conceptualization, S.S., P.L.S., R.F.O-M, T.A.J.G., A.A.T, J.C.G., G.T. and X.Y.; Methodology, M.M., M.V.W, A.A.T, and X.Y.; Validation, P.L.S., and X.Y.; Investigation, D.R., M.M, M.W.V, X.Y, J.C.G., G.T. and A.A.T.; Formal Analysis, D.R., P.L.S., M.M., X.Y, J.C.G., G.T. and T.A.J.G; Resources, R.A.S; Writing-Original Draft, T.A.J.G. and X.Y.; Writing-Review and edit, P.L.S., D.R., R.A.S, R.F.O-M, M.M., M.V.M., A.A.T. J.C.G., G.T. and S.S.; Visualization, P.L.S., D.R. T.AJ.G, J.C.G., G.T. and X.Y; Supervision, S.S and J.W.

## Methods

### Cloning and expression

Residues 46-604 of human ME3 [4] were cloned into pEX-C-His containing a C-terminal TEV protease site and Hexa-His tag. ME3^46-604^ was overexpressed in *E. coli* BL21 (DE3) cells and induced with 0.2 mM IPTG at 16°C for 17 hours.

### Protein purification for structural biology

Frozen cell pellets were resuspended in lysis buffer (20 mM Tris-HCl, pH 8.0, 20 mM Imidazole, pH 8.0, 500 mM NaCl, 3 mM MgCl_2_, 14 mM β-mercaptoethanol (βME), 50 U/ml TurboNuclease and 1X cOmplete Protease Inhibitor) and lysed with a microfluidizer. The whole cell lysate was clarified at 38,400 g for 60 min at 4°C and the supernatant was loaded onto 5 ml HisTrap HP column (GE). The column was washed with 10 column volumes of 10% of buffer B (20 mM Tris-HCl, pH 8.0, 500 mM NaCl, 3 mM MgCl_2_ and 14 mM βME, 400 mM imidazole) and ME3 was eluted with a gradient of 10-100% Buffer B over 20 column volumes (CV). Fractions were pooled and dialyzed overnight at 4°C with TurboTEV (Accelagen) protease (TurboTEV: ME3 = 1:5) in dialysis buffer (20 mM Tris-HCl pH 8.0, 500 mM NaCl, 14 mM βME). Minor residual non-cleaved protein and TurboTEV were removed by applying dialyzed protein to a HisTrap HP column and collecting the flow through and wash. Pooled fractions were concentrated and applied to Superdex S200 (GE Healthcare) size exclusion column (50 mM Tris-HCl pH 8.0, 150 mM NaCl, 1 mM TCEP). In an attempt to remove minor degradation products, the protein was further purified with anion exchange (HiTrap Q, GE healthcare) with limited success. Briefly the protein was desalted into buffer A (25 mM Tris-HCl pH 8.0, 12.5 mM NaCl, 1 mM TCEP) and eluted from the anion exchange column with a gradient elution (12.5mM to 250mM NaCl) over 20 CV. Final pooled eluate was buffer exchanged into storage buffer (50 mM Tris-HCl pH 8.0, 150 mM NaCl, 1 mM TCEP) and concentrated to 1 mg/mL (prep01A) and 11.3 mg/mL (prep01B) for assays and structural determination, respectively.

### Crystallization of ME3 with and without the NADP^+^ cofactor

Initial crystals of ME3 were obtained by sparse matrix screening using a Mosquito crystallization robot (TTP LabTech) at 4°C. The drops consisted of 0.2 μL of Hexa-His tag. ME3^46-604^ (8 mg/mL, 30 mM Tris HCl pH 7.4, 70 mM KCl, 2 mM DTT) and 0.2 μL of reservoir solution. Initial hits were observed in JCSG+ (Molecular Dimensions) B12, PegRx (Hampton Research) F10 and Peg/Ion (Hampton Research) D11.

Data quality crystals were obtained by microseed matrix screening using a Mosquito crystallization robot. The seed stock was prepared using crystals obtained from initial screening, JCSG+ condition B12 (0.2 M Potassium Citrate tribasic, 20% w/v PEG 3350). Crystals were harvested by pipetting the entire 0.4 μL drop into a Hampton seed bead containing 50 μL stabilization buffer and vortexed in 30 second intervals for approximately 4 minutes. Sparse matrix screens were set up using 0.2 μL ME3 (8 mg/mL, Tris HCl pH 7.4, 70 mM KCl, 2 mM DTT), 0.16 μL reservoir solution and 0.04 μL seed stock at 4°C. Singular plates and rods were observed in Peg/Ion condition G5, 0.1 M Ammonium tartrate dibasic pH 7.0, 12% w/v PEG 3350. The final concentration of citrate in the crystal condition was approximately 20 mM.

To obtain a NADP^+^-bound structure of ME3, the citrate-bound ME3 crystals above were soaked overnight in a soaking solution of containing 1 μL of 100mM NADP^+^ and 19 μL of well solution (12% w/v Peg3350, 0.1 M Ammonium tartrate dibasic pH 7.0).

All crystals were cryoprotected by quickly touching crystals into a cryo-protect solution containing 26% PEG 3350, 5% glycerol and 0.2 M ammonium tartrate and flash frozen in liquid nitrogen.

### Structure determination ME3 structures in the presence and absence of NADP^+^

Native diffraction data sets were collected for the structures with and without NADP^+^ to 2.49 Å and 1.94 Å resolution respectively at the Advanced Photon Source (Argonne, IL) beamline 17-ID using a DECTRIS PILATUS 6M detector at a temperature of 100-K and a wavelength of 1.0 Å. Data for all structures were processed and scaled in XDS in the space group P2_1_2_1_2 [32].

The structure of ME3 in the absence of NADP^+^ was solved by Molecular Replacement using Phaser (Phenix) (LLG and TFZ scores of 1313 and 37.2, respectively) using a protomer (monomer) of the previously published structure of human ME2 (PDB ID: 1QR6) as an all atom search model[15, 33]. The asymmetric unit consisted of two molecules arranged in a non-physiological homodimer. The initial model was subjected to rigid body refinement, followed by iterative rounds of building and refinement in Coot and Phenix, respectively [34, 35]. Positive Fo-Fc density was used as a guide to model two molecules of citrate from the crystallization conditions into the allosteric binding site located at the dimer interface. Validation of the final model was performed using composite omit maps calculated in Phenix, and MolProbity. The final structure of ME3 in the absence of NADP^+^ consisted of chain A with density for residues 49-604 and six residues from the C-terminal cleaved TEV-protease site and chain B with density for residues 48-604 and residues corresponding to the full C-terminal linker and cleaved TEV-protease site (respectively). Based on the construct used for crystallography, Hexa-His tag. ME3^46-604^, only 2-3 residues were not observed in the structure. 98.40%, and 1.60% of residues were found in favored and allowed regions of the Ramachandran plots, with no residues in the disallowed region and ^~^99.7% of the residues occupying favorable rotamer conformations.

The NADP^+^-bound structure of ME3 was solved by rigid body refinement (*Phenix.refine*) of the final ME3 apo model described against the ME3 with NADP^+^ data set. Iterative rounds of building (Coot) and refinement (Phenix) were performed. Positive Fo-Fc density was observed for and used to guide the modeling of one molecule of NADP^+^ per monomer. Validation of the final model was performed using composite omit maps calculated in Phenix, MolProbity. The final model consisted of chain A, residues 49-604 and six residues from the cleaved C-terminal TEV-protease site and chain B contains residues 48-604 and two residues form the cleaved C-terminal TEV-protease site. Analysis of the Ramachandran statistics using MolProbity indicated that 97.58%, 2.33%, and 0.09% of the residues are in the favored, allowed and disallowed respectively and ~99% of the residues are in favorable rotamers.

In accordance with the literature precedence, the analysis of ME3 in this paper was described using ME2 sequence numbering [9, 20]. Based on this new numbering system, ME3 crystal structures contain residues 23-582 with two breaks in the numbering due to deletions of 1 and 2 residues after residues 354 and 370, respectively (Figure S2).

### Grid preparation and data acquisition.

3 μL of 0.8-1.0 mg/ml purified ME3 with/without Citrate or NADP complex was applied to the plasma-cleaned (Gatan Solarus) Quantifoil 1.2/1.3 UltraAuFoil holey gold grid, and subsequently vitrified using a Vitrobot Mark IV (FEI Company). Cryo grids were loaded into a Titan Krios transmission electron microscope (ThermoFisher Scientific) with a post-column Gatan Image Filter (GIF) operating in nanoprobe at 300 keV with a Gatan K2 Summit direct electron detector and an energy filter slit width of 20 eV. Images were recorded with Leginon in counting mode with a pixel size of 1.04 Å and a nominal defocus range of −2.4 to −1 μm. Data were collected with a dose rate of 6.9 electrons per physical pixel per second, and images were recorded with a 6s exposure and 200 ms subframes (30 total frames) corresponding to a total dose of 44.9 electrons per Å^2^. All details corresponding to individual datasets are summarized in Table 1.

### Electron microscopy data processing.

Dose-fractioned movies were gain-corrected, and beam-induced motion correction using MotionCor2 [36] with the dose-weighting option. The ME3 particles were automatically picked from the dose-weighted, motion corrected average images using Relion 3.0 [37]. CTF parameters were determined by Gctf [38]. Particles were then extracted using Relion 3.0 with a box size of 200 pixels. The 2D, 3D classification and refinement were performed with Relion 3.0. Two rounds of 2D classification and one round of 3D classification were performed to select the homogenous particles. Homogenous particles were then submitted to 3D auto-refinement with the D2 symmetry imposed. All 3D classifications and 3D refinements were started from a 60 Å low-pass filtered version of an ab initio map generated with Relion 3.0. Per-particle CTF estimation was refined using the program Relion 3.0 and followed by one round of 3D auto-refinement. All resolutions were estimated by applying a soft mask around the protein complex density and based on the gold-standard (two halves of data refined independently) FSC = 0.143 criterion. Prior to visualization, all density maps were sharpened by applying different negative temperature factors using automated procedures, along with the half maps, were used for model building. Local resolution was determined using ResMap [39] (Figure S6).

### Model building and refinement

The initial template of the human ME3 was derived from a homology-based model calculated by SWISS-MODEL [40]. The model was docked into the EM density map using Chimera [41] and followed by manual adjustment using COOT [42]. Note that the apo EM density around the peripheral loop regions at Domain C was not sufficient to build the model. Each model was independently subjected to global refinement and minimization in real space using the module *phenix.real_space_refine* in PHENIX [43] against separate EM half-maps with default parameters. The model was refined into a working half-map, and improvement of the model was monitored using the free half map. The geometry parameters of the final models were validated in COOT and using MolProbity [44] and EMRinger [45]. These refinements were performed iteratively until no further improvements were observed. The final refinement statistics were provided in Table 2. Model overfitting was evaluated through its refinement against one cryo-EM half map. FSC curves were calculated between the resulting model and the working half map as well as between the resulting model and the free half and full maps for cross-validation (Figure S6). Figures were produced using PyMOL [46] and Chimera.

### Malic enzyme activity assays

ME2 and ME3 activity was measured by following the absorbance of NADH and NADPH [17]. Buffers containing 50 mM TRIS (pH 7.5), 10 mM MgCl_2_ and either no additive, 5 mM citrate, or 5 mM fumarate were prepared. All three buffers were then titrated to a pH of 7.5 using NaOH. For ME3 kinetic assays, 10nM ME3 enzyme, 10 mM Malate and 0.3 mM NADP were added to each buffer to a final reaction volume of 1 mL. For ME2 kinetic assays, 100 nM ME2 enzyme, 10 mM Malate and 0.3 mM NAD were added to each buffer to a final reaction volume of 1mL. The kinetic assays were performed by adding 1 mL sample to a Thermo Scientific NanoDrop One^C^ Microvolume UV-Vis Spectrophotometer and following A_340_ (absorbance of NAD(P)H)) for 5 minutes. The enzyme reagent was added to the sample immediately prior to adding the sample to the spectrophotometer. Each sample was run in n=3 biological replicates using a 10 mm path-length quartz cuvette.

ME3 cofactor preference was determined using a coupled assay with Diaphorase. ME3 activity was monitored in real time by following the formation of Resorufin using BMG ClarioStar spectrophotometer using excitation wavelength of 545 nm and emission at 600 nm. Reaction mixture contained 50 mM Tris, pH 7.5; 50 mM NaCl; 0.01% Triton X 100, 1% DMSO, 10 mM MgCl_2_, 10 mM Malate, 0.015 mg/ml diaphorase and 50 μM resazurin and the indicated concentrations of either NADP^+^ or NAD^+^. The assay was initiated by addition of 1nM of ME3 to the reaction mixture, to a final volume of 20 μL.

