## Supporting Data for "Structures of human Malic Enzyme 3"

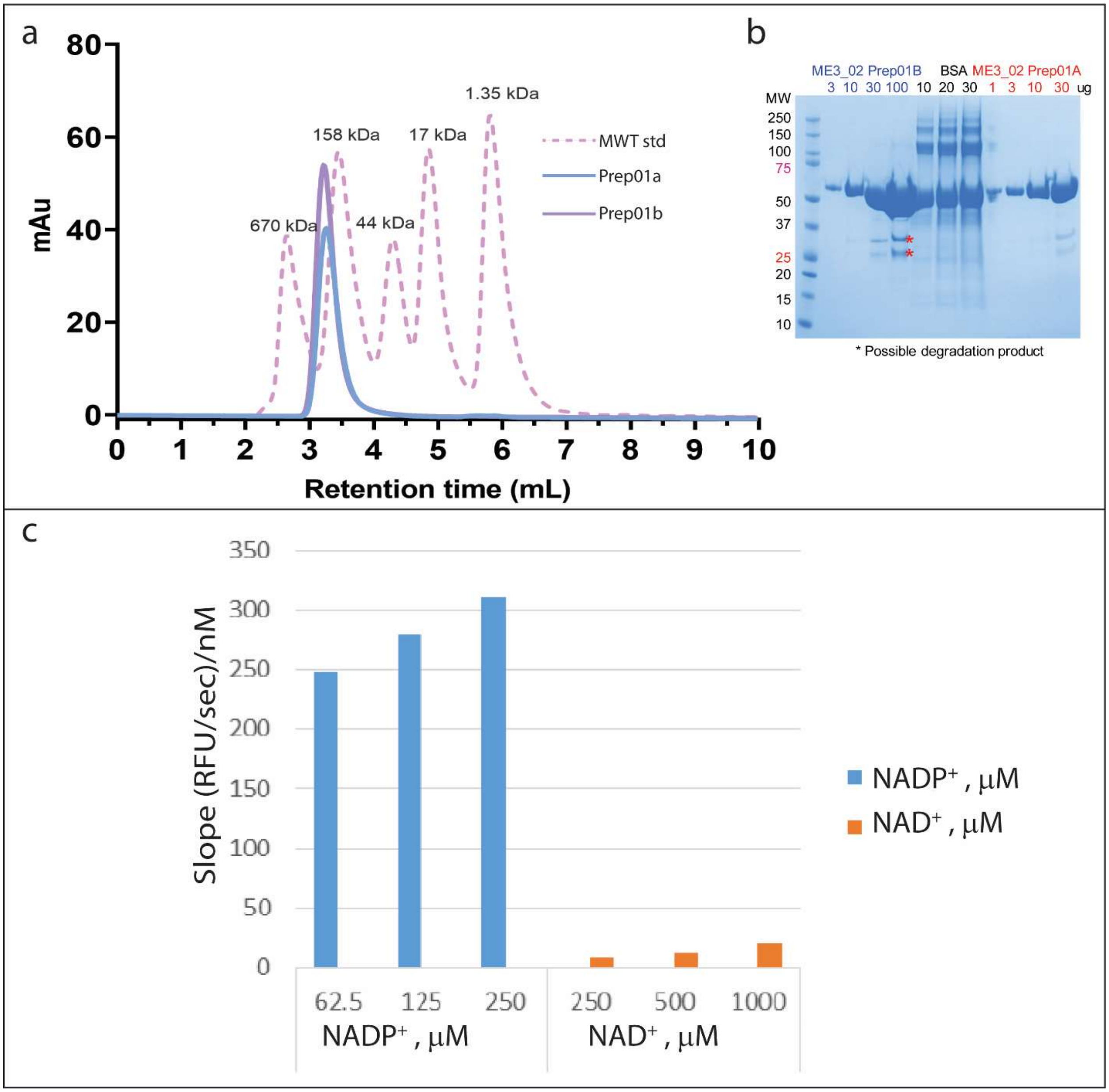


**Figure S1.** Biophysical and biochemical characterization of purified ME3. **(a)** Analytical size exclusion chromatogram for the purified of ME3, Prep01A (in blue) and Prep01B (in purple), used in assays and structural biology, respectively. The molecular weight standard is shown as pink dashed lines and the molecular weights are indicated in black. **(b)** Coomassie stained SDS-PAGE gel of purified ME3 used in this paper. Red stars indicate possible degradation. **(c)** The activity of ME3 in the presence of $NADP^+$ (blue) and $NAD^+$ (orange) is represented by a bar graph.

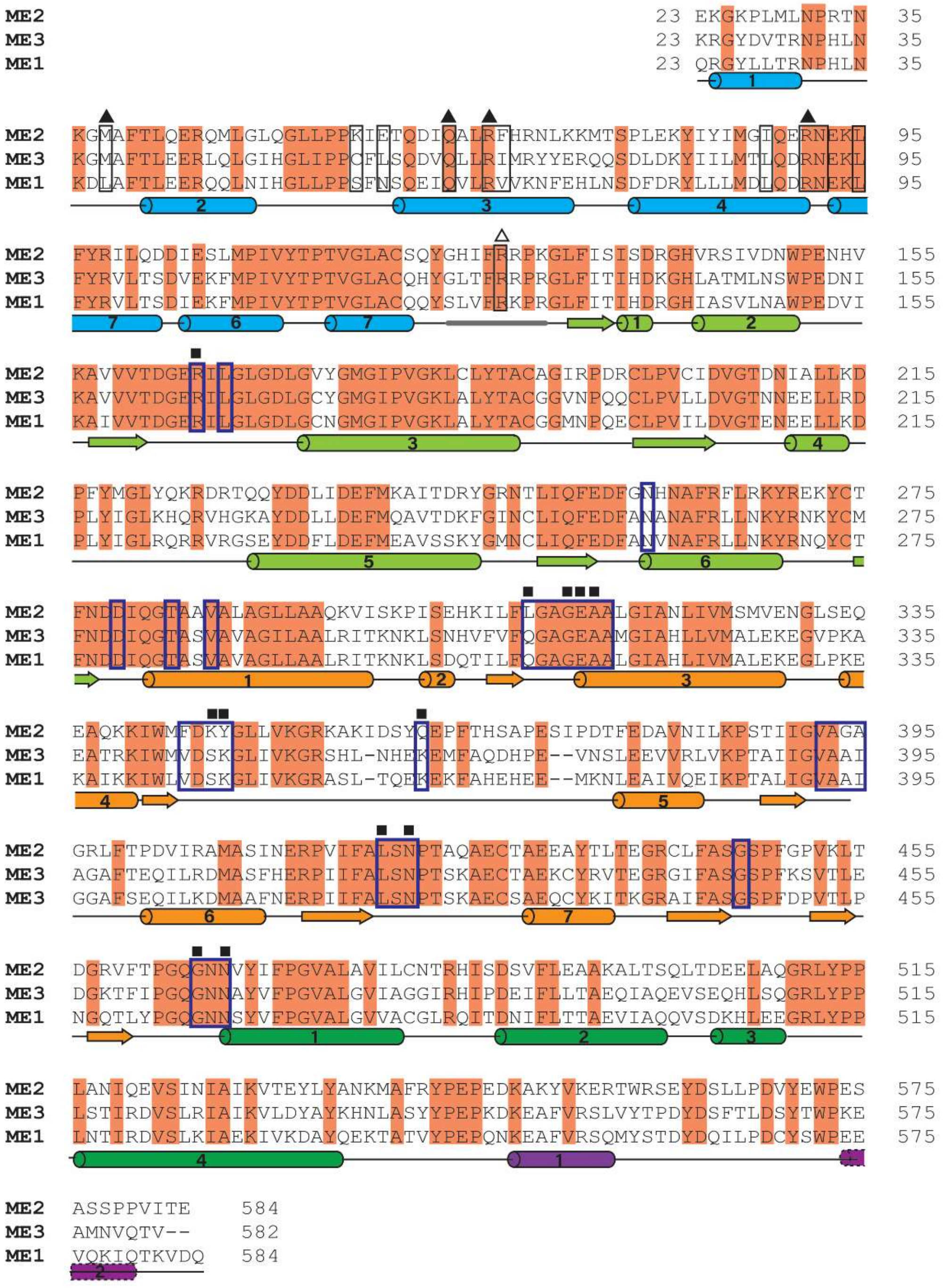


**Figure S2.** Structure-based sequence alignment of the human MEs truncated to reflect the crystallography constructs and numbered according to the ME2 human isoform. Secondary structure elements are shown according to the crystal and EM structures of the human ME3 and color coded as individual domains. Loop AB is shown as thick gray line. Conserved residues are highlighted in orange and residues interacting with citrate or $NADP^+$ are highlighted with black or thick blue open boxes, respectively. Key residues are indicated as triangle or square above the sequence for the interactions with citrate and $NADP^+$, respectively, with the filled ones from one protomer and the open ones from the neighbor protomer in the asymmetric unit.

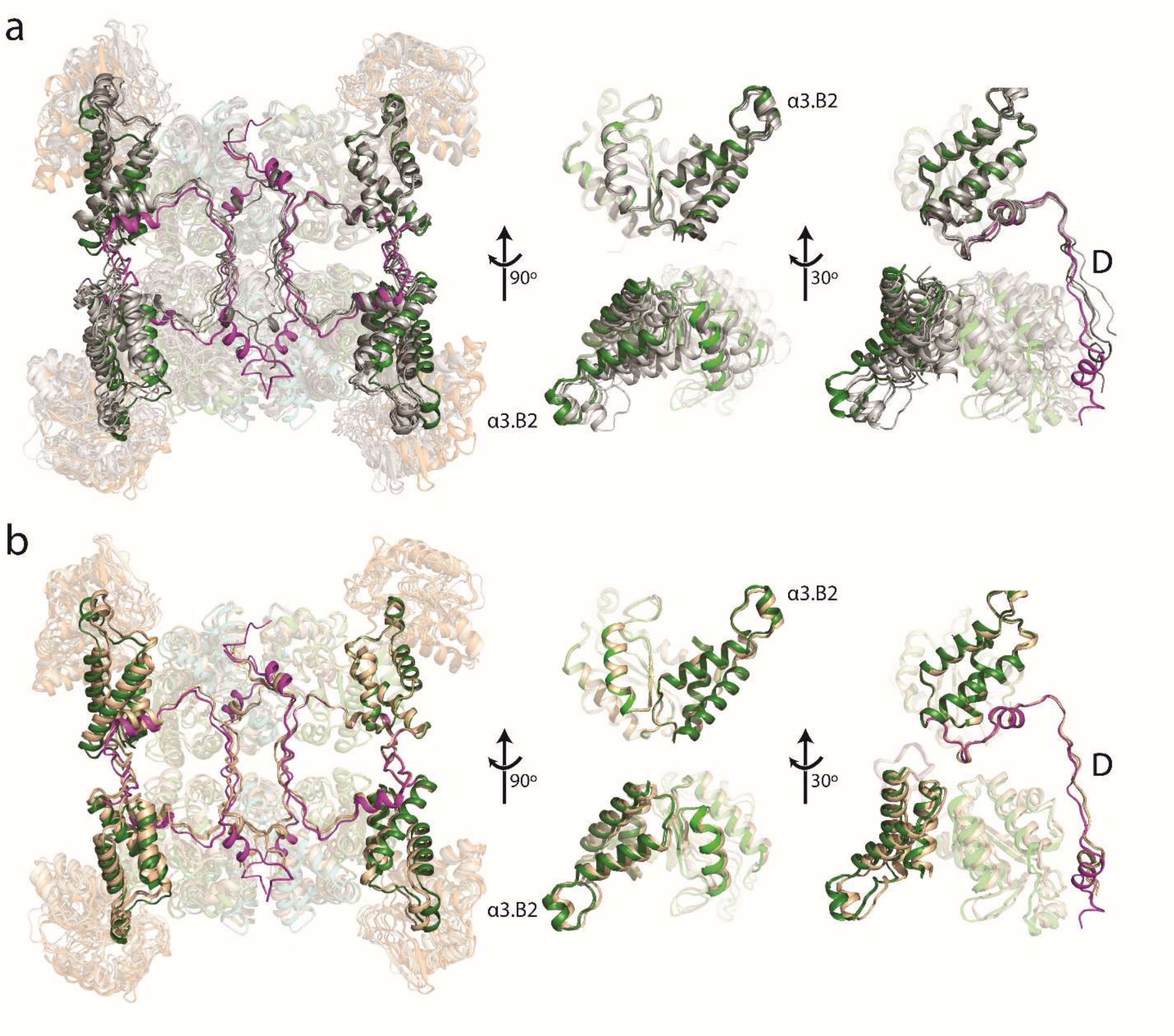


**Figure S3.** Structural superposition of the Xray structures of MEs **(a)**, and Xray and EM structures of ME3 **(b)** in the tetrameric complexes. The Xray tetrameric complex was generated by crystallographic symmetry operation. The structures were superimposed using the corresponding domain B2 from one of the protomer in the tetrameric complex. The Xray structures of ME3 were color coded as individual domains. The Xray structures of ME1 and ME2 are in dark gray and white, respectively. The EM structures of ME3 are in wheat. Domains B2 and D are highlighted. The helix α3 from domain B are shown as the reference.

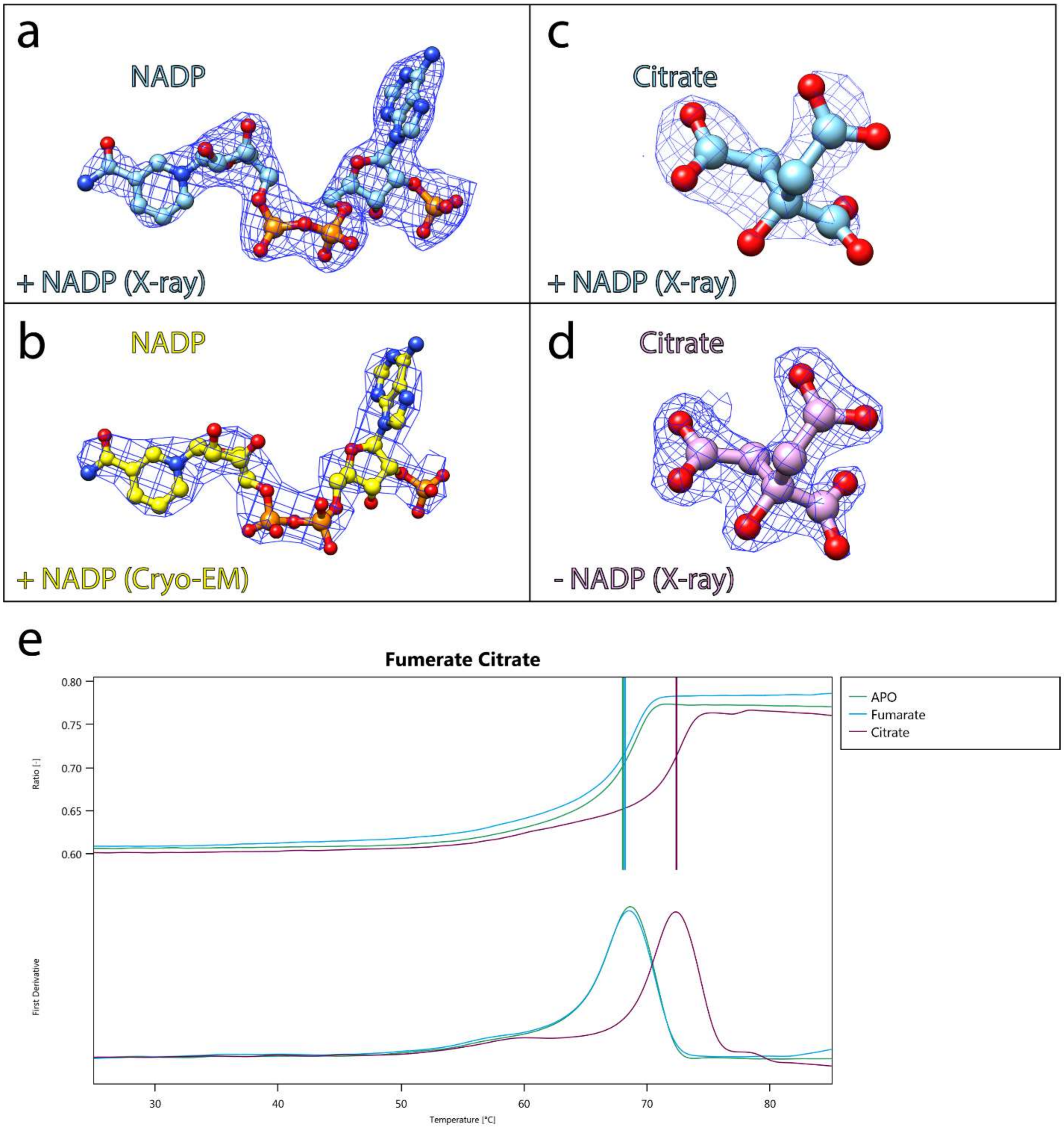


**Figure S4**. Comparison of map quality of bound NADP⁺ (left panels) and citrate (right panels) molecules in the ME3 cryo-EM and X-ray structures. Electron density is shown for **(a)** NADP⁺ and **(c)** citrate bound to the crystal structure of ME3 solved to 2.49 Å -resolution. **(b)** Electron density for NADP⁺ molecule bound in the cryo-EM structures solved in the presence of NADP⁺ to 2.77 Å-resolution. **(d)** Map quality of citrate bound to the X-ray crystal structure solved in the absence of NADP⁺ to 1.94 Å-resolution is shown. **(e)** NanoDSF thermal melting traces for ME3 in the presence of Fumarate (blue), citrate (purple) and in the absence of ligands, Apo (blue).

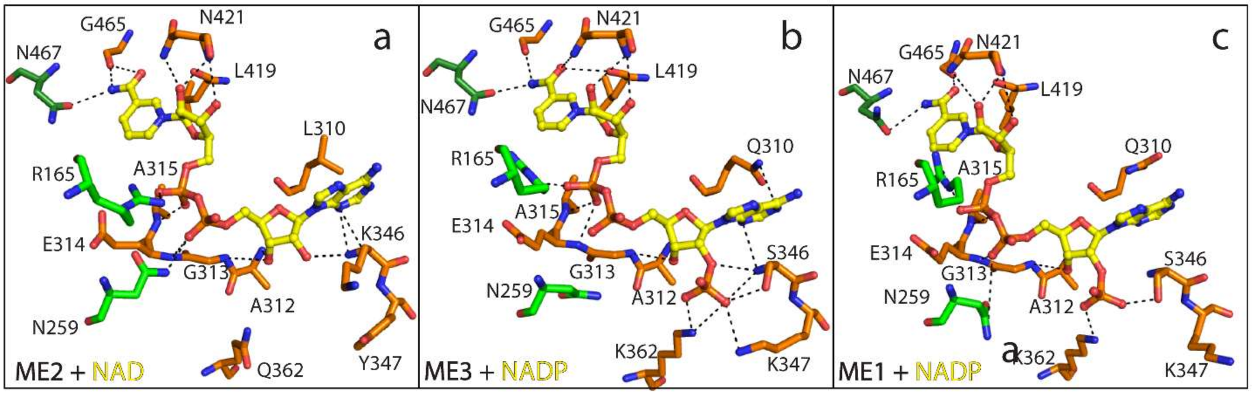


**Figure S5.** The NAD(P)$^+$ binding pocket of ME2 **(a)** ME3 **(b)** and ME1 **(c)** with the cofactor in ball and stick representation and colored yellow ball. The residues are colored by domains with Domain B in green and Domain C in orange. The ligand binding interactions are indicated by dash lines.

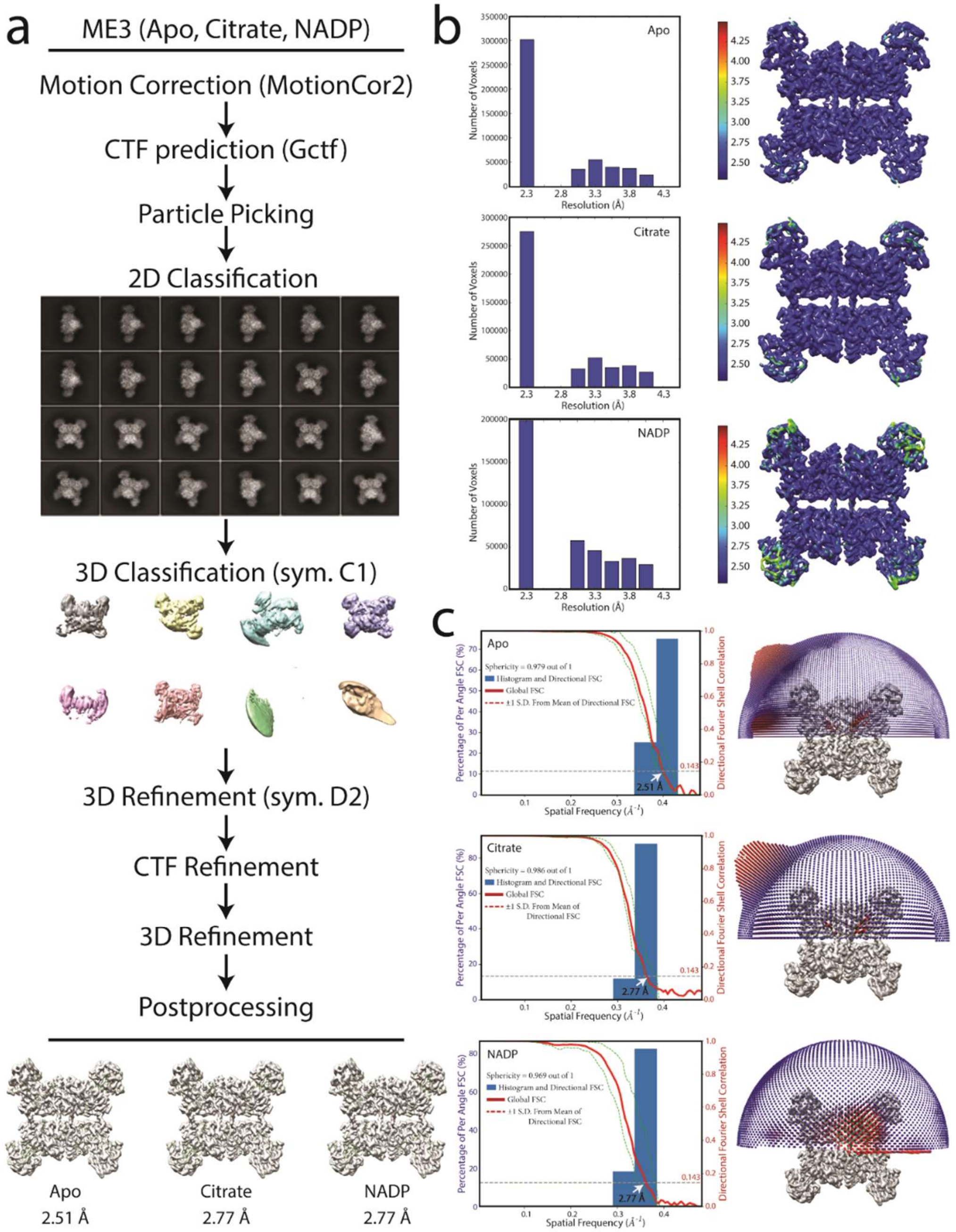


**Figure S6.** Cryo-EM analysis of ME3 complex. **(a)** Flow chart of the cryo-EM data processing procedure. Details can be found in the Methods. **(b)** Local resolution of the maps estimated using the ResMap program and colored as indicated. **(c)** Gold standard FSC curves of the structures and Angular orientation distribution of the particles used in the final reconstruction. The particle distribution is indicated by different color shades.

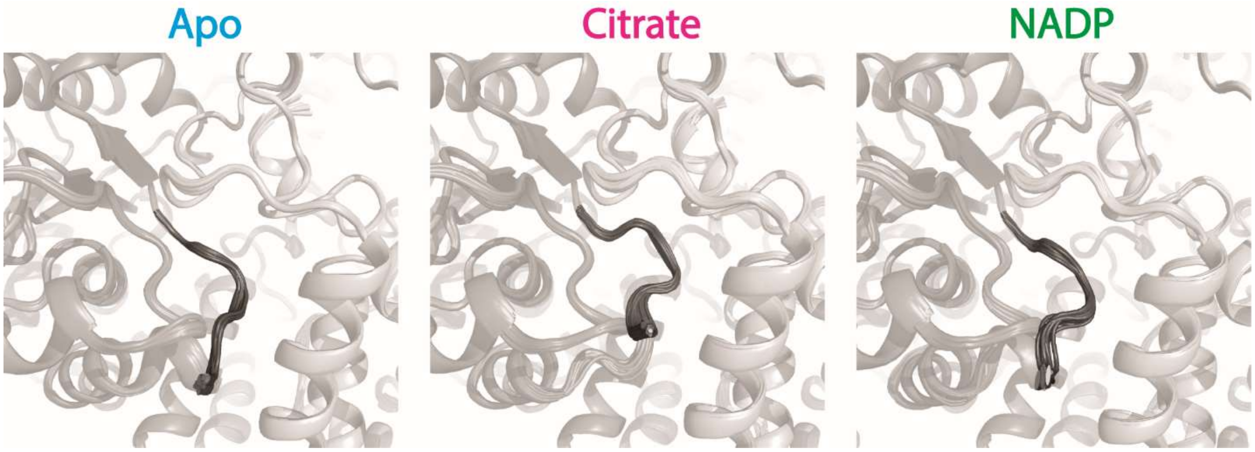


**Figure S7.** Dynamics of Loop AB in ME3 cryoEM structures. Overlay of 50 refined structures in Rosetta. Loop AB was colored in solid gray. The ensemble at Loop AB displays a higher degree of flexibility in the Apo, Citrate and NADP binding states.

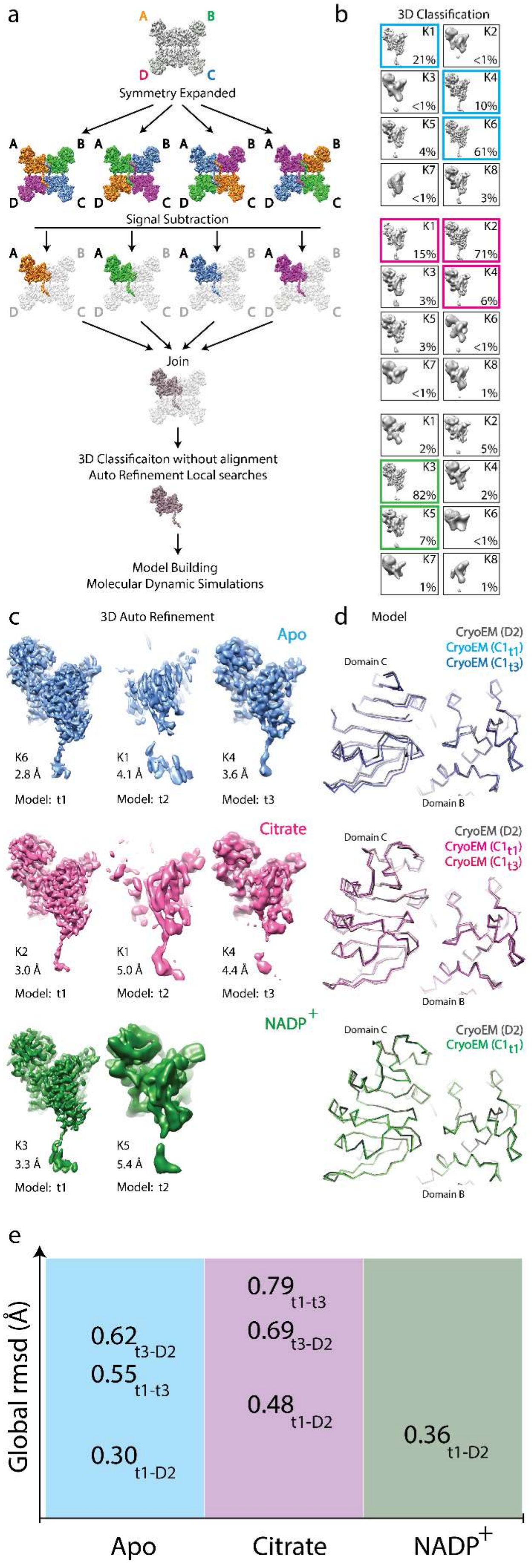
a
A
B
D
C
Symmetry Expanded
Signal Subtraction
Join
3D Classificaiton without alignment
Auto Refinement Local searches
Model Building
Molecular Dynamic Simulations
b
3D Classification
K1 21%
K2 <1%
K3 <1%
K4 10%
K5 4%
K6 61%
K7 <1%
K8 3%
K1 15%
K2 71%
K3 3%
K4 6%
K5 3%
K6 <1%
K7 <1%
K8 1%
K1 2%
K2 5%
K3 82%
K4 2%
K5 7%
K6 <1%
K7 1%
K8 1%
c
3D Auto Refinement
Apo
K6 2.8 Å
Model: t1
K1 4.1 Å
Model: t2
K4 3.6 Å
Model: t3
Citrate
K2 3.0 Å
Model: t1
K1 5.0 Å
Model: t2
K4 4.4 Å
Model: t3
NADP+
K3 3.3 Å
Model: t1
K5 5.4 Å
Model: t2
d
Model
CryoEM (D2)
CryoEM (C1t1)
CryoEM (C1t3)
Domain C
Domain B
e
Global rmsd (Å)
0.62 t3-D2
0.55 t1-t3
0.30 t1-D2
0.79 t1-t3
0.69 t3-D2
0.48 t1-D2
0.36 t1-D2
Apo
Citrate
NADP+

**Figure S8.** Subunit conformational variation within individual ME3 tetramers resolved by cryo-EM. **(a)** Flow chart of symmetry expansion and followed by density subtraction to investigate dynamic features of ME3. D2 symmetry expansion resulted in 4 set of particle images, with each of the 4 subunits aligned to the same orientation. Individual protomers were color coded related to their original corresponding positions (A, B, C, and D). Protomer signals in positions B, C, and D were subtracted from the respective symmetry-expanded images. The subparticle images containing only the signal of a single protomer and were joined and subjected to 3D focused classification while keeping all of the orientations fixed at the values determined in the refinement of the original maps. **(b)** This method of image analysis was used to separate 8 classes of ME3. The number of subparticles within each class is specified. **(c)** Localized reconstruction from the major converged classes of subunits. **(d)** Superposition of the models derived from the cryo-EM maps of D2 symmetry or signal subtractions with C1 symmetry. **(e)** Global r.m.s.d.s of structures solved from 3D classification performed without alignment or symmetry against structures with D2 symmetry.

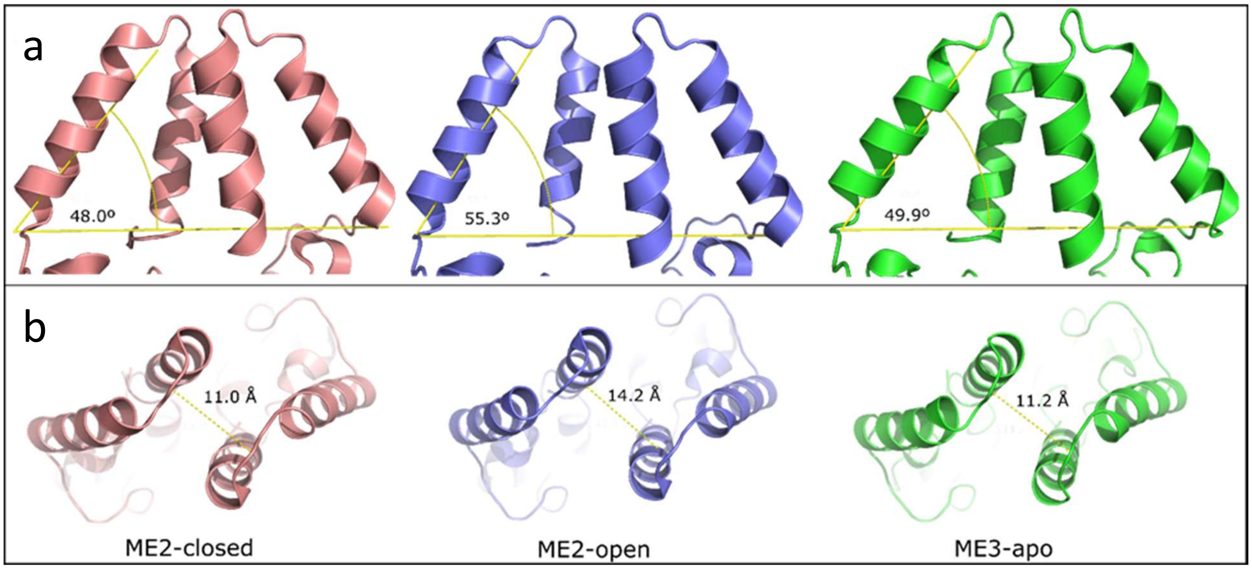


**Figure S9**: Domain A dimer interface comparison. **(a)** The angle formed between the domain A of each monomer and **(b)** Distance between Thr76 (Gln in ME3) between monomers for ME2-closed (dark pink), ME2-open (dark blue), and ME3 in apo form (green). The interface distances are 11.0, 11.2, and 14.2 Angstroms for ME2-closed, ME3, and ME2-open structures respectively. The angle between the helices at the interface is 48.0, 49.9, and 55.3 degrees for ME2-closed, ME3, and ME2-open structures respectively. The angle taken as reference is formed by Thr60 (Ser in ME3) at the base of helix I, Thr60 in the other monomer, and MSE75 (Gln in ME3) at the top of helix I.

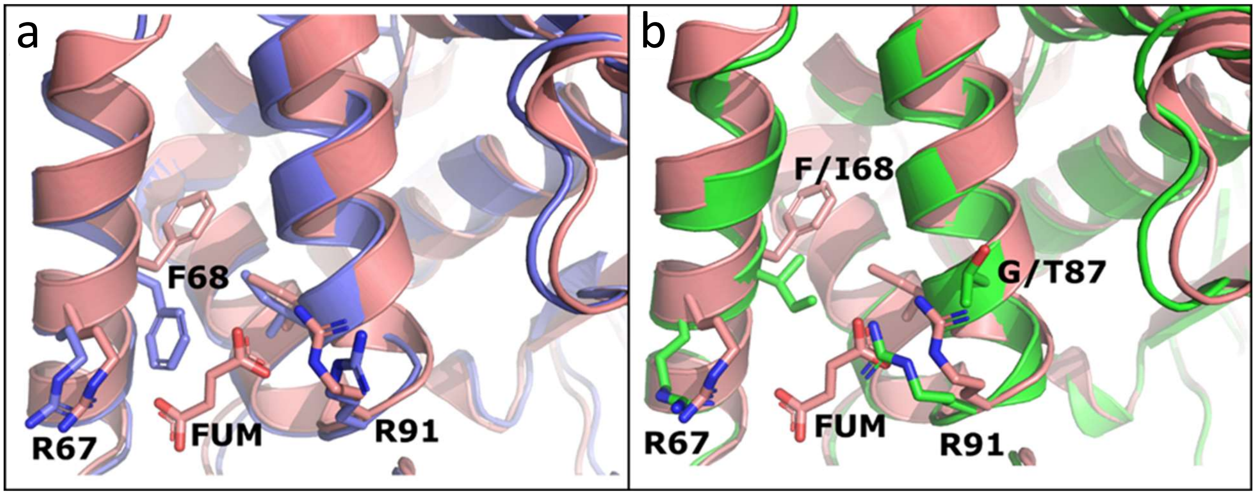


**Figure S10**: Domain A superposition ME2 open (blue), ME2 closed (pink), and ME3 (green). **(a)** Fully active ME2 shows a conformational rearrangement of domain A. A tilt towards the dimer geometrical center together with an inward rotamer switch of Phe68 in the active conformation (pink) which is also compatible with fumarate binding. **(b)** ME3 lacks Phe68 (Ile) and shows a closer arrangement to ME2 in its closed form. Also, Thr87 in ME3 (Gly in ME2) prevents Arg91 to adopt a conformation compatible with fumarate binding.

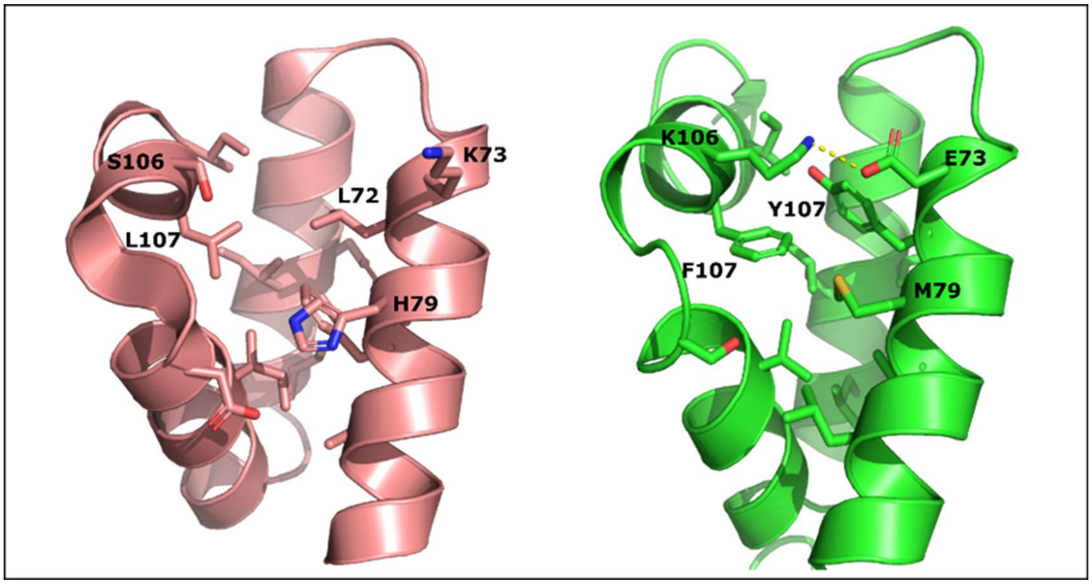


**Figure S11: Domain A packing**. Comparison between residues forming the core of domain A in ME2 (dark pink) and ME3 (green). ME3 shows a more compact core due to the contribution of Tyr72 and Phe107 (both leucine in ME2). Also, ME3 shows an ionic interaction between Glu73 and Lys106 which is not seen in ME2 (Lys73-Ser106).

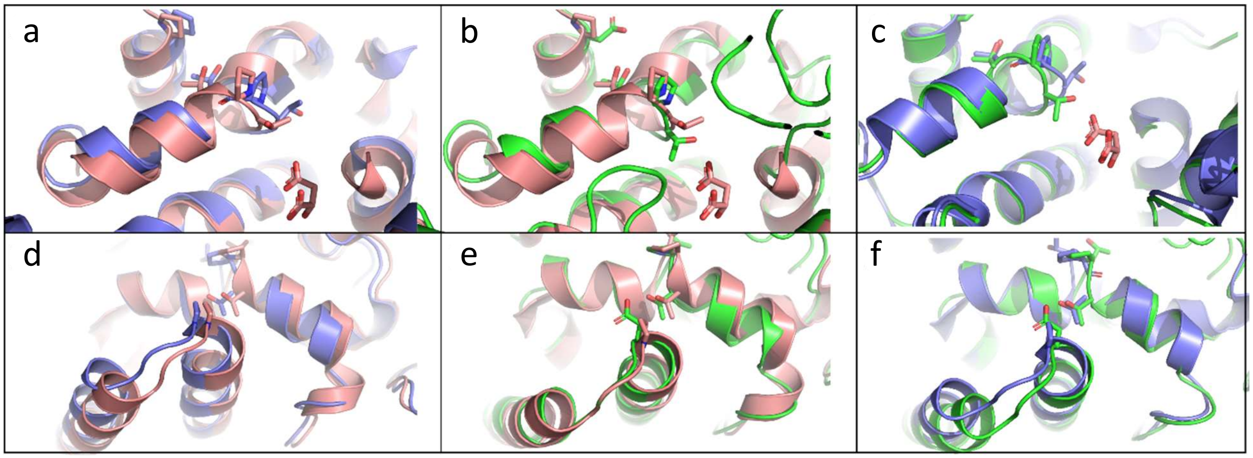


**Figure S12: Effect of domain A conformational change at substrate binding pocket**. **(a,b,c)**: Substrate binding site detail in the superposition of ME2 open (dark blue), ME2 closed (pink), and ME3 apo (green) Domain A. **(d,e,f)**: Dimer interface detail of Domain A helices. Conformational change of the two interface helices promotes a change in the perpendicular domain A helix through a mechanical push of Pro78 to Thr115. This change occurring at the helix kink formed by Thr113 and Pro114 leaves Thr113 directly facing the substrate. Domain A ME3 superposition to the ME2 closed form shows that ME3 in the apo form already has a similar arrangement without the binding of cofactors or allosteric modulators.

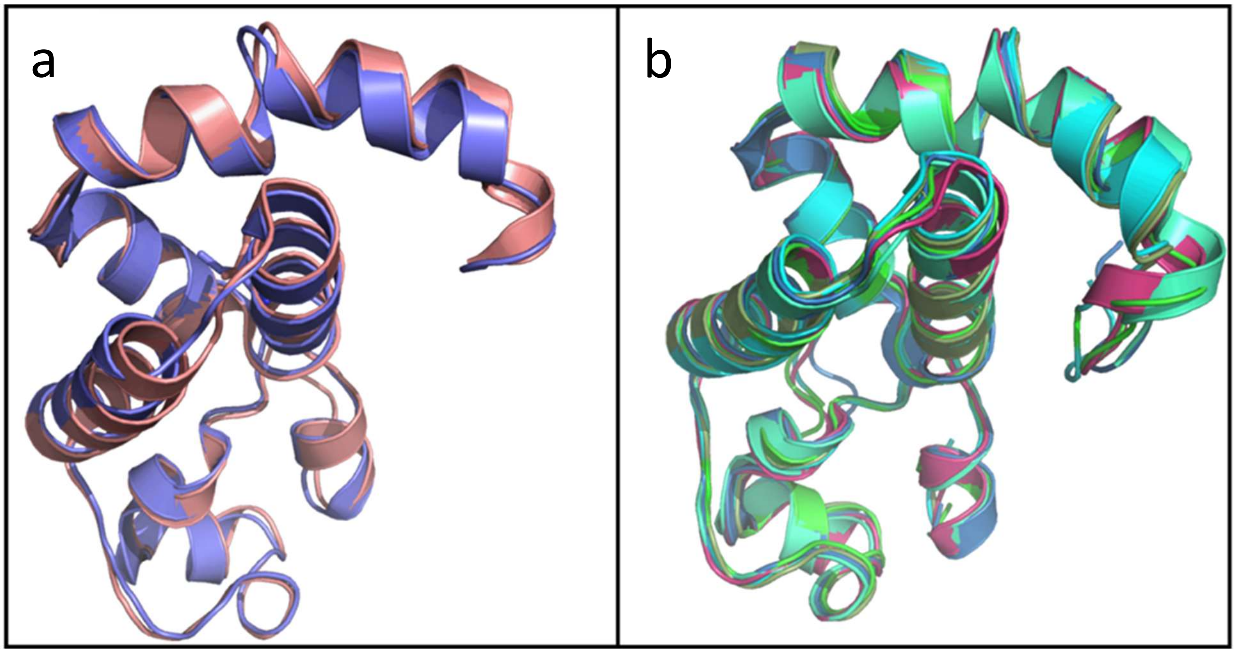


**Figure S13:** Comparison between ME2 and ME1-ME3 superposition of Domain A. **(a)** Superposition of domain A in ME2 closed (dark blue) and open (dark pink) forms (PDB IDs: 1pj2, 1qr6) unveils a conformational change in the interface helices leading to a change in the kink of the transversal helix (top of each panel). **(b)** Superposition of ME1 and ME3 structures (ME3 apo, ME3 bound to citrate, and ME3 bound to citrate and NADP+ from this work, human ME1 in apo form (PDB ID: 3wja) and pigeon liver ME1 in closed form (PDB ID: 1gq2). ME3 and ME1 show a very close arrangement of the domain regardless of the bound state.

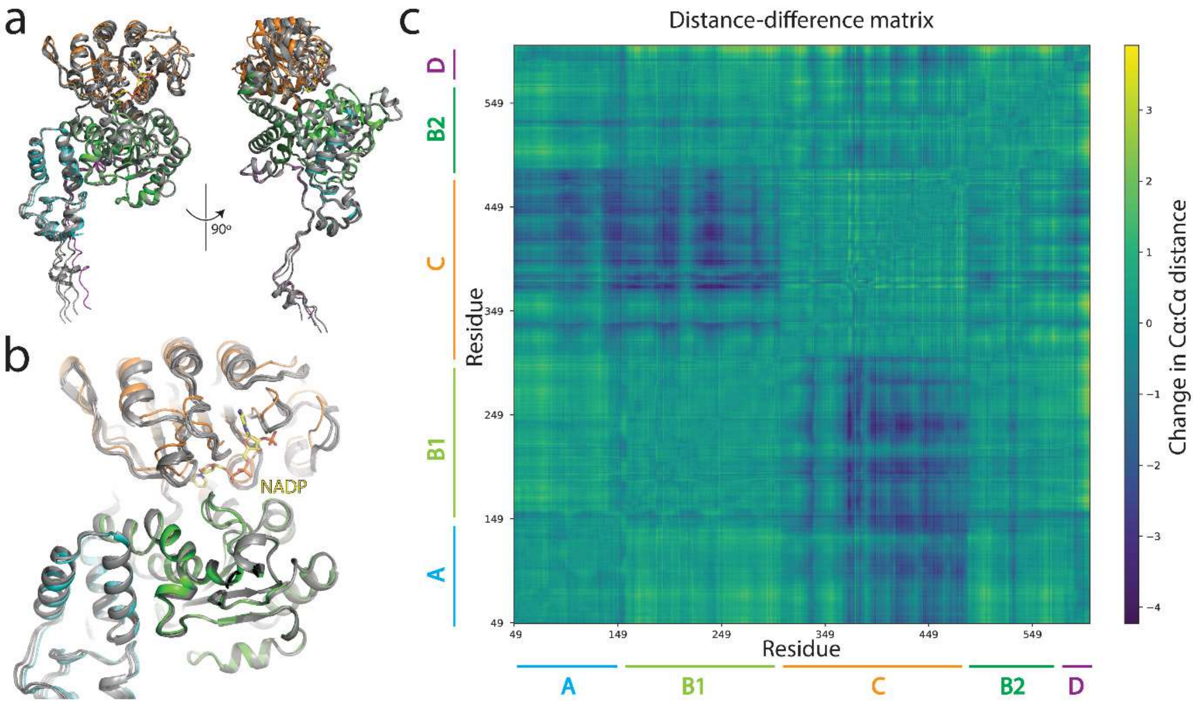

**Figure S14. (a)** Overlay of 5 AlphaFold2 predicted models with the NADP bound cryo-EM ME3 structure. **(b)** Closed view at the substrate binding site. Alphafold2 models were colored gray and NADP-bound Cryo-EM ME3 structure followed the color code as in Figure 1c. **(c)** Distance-difference matrix was calculated using the ME3 NADP-bound cryo-EM structure and Alphafold2 model. Individual domains were highlighted.

| Sequence Identity % / RMSD (Å) | Global | A | B1 | C | B2 | D |
|---|---|---|---|---|---|---|
| **ME1 - ME2** | 55.5 / 1.21 | 54.6 / 0.74 | 66.7 / 0.58 | 53.4 / 1.11 | 47.2 / 0.92 | 44.1 / 0.99 |
| **ME1 - ME3** | 71.6 / 1.18 | 72.0 / 1.13 | 81.0 / 0.56 | 73.7 / 0.83 | 68.6 / 0.74 | 54.8 / 1.74 |
| **ME2 – ME3** | 56.1 / 1.50 | 57.9 / 1.10 | 66.0 / 0.75 | 53.4 / 1.11 | 52.2 / 0.90 | 44.1 / 1.32 |
| **ME3 – $ME3_{EM}$** | 100.0 / 1.12 | 100.0 / 0.88 | 100.0 / 0.51 | 100.0 / 0.71 | 100.0 / 0.35 | 100.0 / 1.24 |

**Table S1**. Amino acid identity and average main chain RMSD values (in Å) between protomers of human ME1 (PDB ID: 3WJA), ME2 (PDB ID: 1PJ3), NADP bound ME3 X-ray and EM structures (this study) are indicated for each domain and across total protein length. The RMSD values were calculated using Superpose in CCP4.

| | ME | Organism | PDB Code | Method | Cofactor | Substrate | Cation | Activator/Ligand | Form |
|---|---|---|---|---|---|---|---|---|---|
| **1** | c-NADP-ME | Human | 3WJA | Xray | | | | | Open |
| **2** | c-NADP-ME | Columba livia | 1GQ2 | Xray | NAP | OXL | MN | | Closed |
| **3** | m-NAD(P)-ME | Human | 1QR6 | Xray | NAD | | | | Open |
| **4** | m-NAD(P)-ME | Human | 1PJL | Xray | NAD | | LU | | Open |
| **5** | m-NAD(P)-ME | Human | 1DO8 | Xray | NAD | OXL | MN | | Closed |
| **6** | m-NAD(P)-ME | Human | 1EFK | Xray | NAD | MAK | MN | | Closed |
| **7** | m-NAD(P)-ME | Human | 1EFL | Xray | NAD | TTN | MN | | Closed |
| **8** | m-NAD(P)-ME | Human | 1PJ3 | Xray | NAD | PYR | MN | FUM | Closed |
| **9** | m-NAD(P)-ME | Human | 1PJ2 | Xray | NADH | LMR | MN | FUM | Closed |
| **10** | m-NAD(P)-ME | Human | 1GZ4 | Xray | ATP | TTN | MN | FUM | Closed |
| **11** | m-NAD(P)-ME | Human | 1PJ4 | Xray | ATP | MLT | MN | FUM | Closed |
| **12** | m-NAD(P)-ME | Human | 1GZ3 | Xray | ATP | OXL | MN | FUM | Closed |
| **13** | m-NAD(P)-ME | Ascaris suum | 1LLQ | Xray | NAD | | | | Open |
| **14** | m-NAD(P)-ME | Ascaris suum | 1O0S | Xray | NADH | TTN | | | Open |
| **15** | m-NADP-ME | Human | This study | Xray | NAP | | | CIT | Open |
| **16** | m-NADP-ME | Human | This study | Xray | | | | CIT | Open |
| **17** | m-NADP-ME | Human | This study | EM | | | | CIT | Open |
| **18** | m-NADP-ME | Human | This study | EM | NAP | | | | Open |
| **19** | m-NADP-ME | Human | This study | EM | | | | | Open |

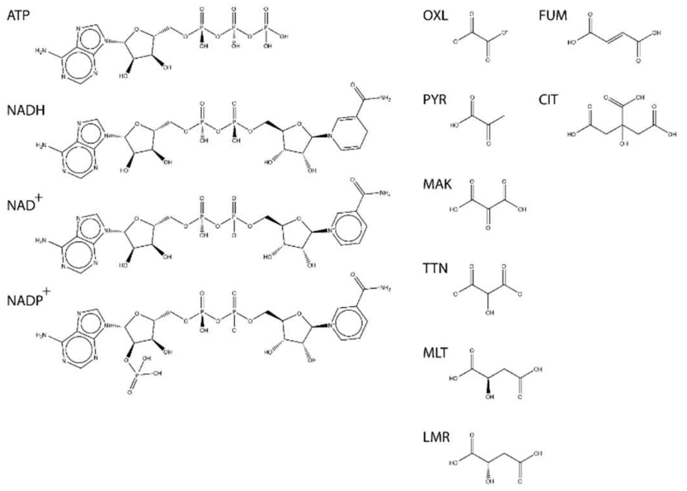


**OXL** OXALATE ION
**MAK** ALPHA-KETOMALONIC ACID
**TTN** TARTRONATE
**LMR** (2S)-2-hydroxybutanedioic acid
**PYR** PYRUVIC ACID
**MLT** D-MALATE
**FUM** FUMARIC ACID
**CIT** CITRIC ACID
**NAD+** NICOTINAMIDE-ADENINE-DINUCLEOTIDE
**NADH** 1,4-DIHYDRONICOTINAMIDE ADENINE DINUCLEOTIDE
**NADP+** NADP NICOTINAMIDE-ADENINE-DINUCLEOTIDE PHOSPHATE

**Table S2.** (a) Details of Malic enzymes structures, published and in this paper, including the structure determination methods, ligands bound, and the active site form. (b) Chemical structures of ligands observed in malic enzyme structures.
